# Low photorespiratory capacity is sufficient to block the induction of photosynthetic capacity in high light

**DOI:** 10.1101/2021.01.20.427534

**Authors:** Christopher R. Baker, Jean Christophe Cocurrun, Ana Paula Alonso, Krishna K. Niyogi

## Abstract

The induction of high photosynthetic capacity in high light (HL) is a common response among many herbaceous dicot plants, however, the signals that control this response remain largely unknown. Here, multiple independent lines of evidence are presented in support of the conclusion that low photorespiratory capacity acts a negative signal to limit photosynthetic capacity acclimation in HL in *Arabidopsis thaliana*. Using a panel of natural accessions, primary nitrogen (N) assimilation and photorespiration rates early after a shift to growth in HL, as well as activities for key enzymes in these pathways, were shown to positively correlate with the magnitude of the subsequent induction of photosynthetic capacity, which occurred several days later. Time-resolved metabolomic data during acclimation to HL were collected using a strongly acclimating ecotype and a weakly acclimating ecotype, revealing in greater detail the differences in N assimilation, photorespiration, and triose-phosphate utilization pathways underlying efficient photosynthetic capacity acclimation. When shifted into HL growth conditions under non-photorespiratory conditions, weakly acclimating ecotypes and even photorespiratory mutants gained the ability to strongly induce high photosynthetic capacity in HL. Thus, a negative, photorespiration-dependent signal early in the HL shift appears to block photosynthetic capacity acclimation in accessions with low photorespiratory capacity, whereas accessions with high photorespiratory capacity are licensed to increase photosynthetic capacity.

## Introduction

Photosynthetic organisms tune photosynthetic capacity (i.e., the CO_2_- and light-saturated maximum photosynthetic rate) in response to environmental and developmental signals (Björkman and Holmgren, 1963; Evans and Poorter, 2001; Athanasiou et al., 2010; Demmig-Adams et al., 2017). For instance, many herbaceous dicot plants increase photosynthetic capacity in response to growth in high light (HL) relative to growth in low light (LL) (Murchie and Horton, 1997; Björkman, 1981). Increasing photosynthetic capacity in crops has been proposed as a means to increase agricultural yields in this century (Long et al., 2015; Parry et al., 2011; Driever et al., 2014). An understanding of how endogenous signaling and metabolic acclimation responses tune photosynthetic capacity in response to the growth environment will accelerate and support engineering approaches to increase photosynthetic capacity.

The acclimation of photosynthetic capacity to HL is part of the suite of acclimation responses that maintain redox poise of the photosynthetic electron transport (PET) chain in HL (Anderson and Osmond, 1987; Savitch et al., 2000; Park et al., 1996; Demmig-Adams et al., 2017; Herrmann et al., 2019). The PET chain is reduced in HL as excess excitation energy exceeds the capacity of downstream reactions to utilize the end products of PET (Li et al., 2009). q_L_, a chlorophyll fluorescence parameter that measures the oxidation state of the photosystem II (PSII) electron acceptor site Q_A,_ can be used to quantify the redox state of the PET chain (Kramer et al., 2004). To increase q_L_ under HL growth (i.e., to oxidize the PET chain), photosynthetic organisms can decrease light absorption and/or divert a greater fraction of absorbed light energy into non-photochemical energy dissipation mechanisms (Bailey et al., 2001; Anderson et al., 1988; Bailey et al., 2004). They can also increase utilization of the products of PET thereby mitigating end-product inhibition of PET in HL (Hald et al., 2008; Pammenter et al., 1993; Foyer et al., 2003). On a longer timescale, increased photosynthetic capacity via greater resource allocation to building the components of PET and the Calvin-Benson-Bassham (CBB) cycle will also increase q_L_ in HL (Walters et al., 2003; Evans and Poorter, 2001; Björkman, 1981). Thus, the acclimation of photosynthetic capacity in HL can be thought of both as a means to balance photosynthetic energy production with sink utilization and as a photoprotective response necessary in excess light environments.

Photosynthetic capacity can be estimated using several different approaches including: (1) chlorophyll fluorescence-derived estimation of the light-saturated, maximum electron transport rate (ETR_max_), (2) the CO_2_- and light-saturated, maximum oxygen evolution rate (P_max_), (3) the CO_2_- and light-saturated, maximum CO_2_ assimilation rate (A_max_), and (4) the use of a CO_2_ assimilation curve (an A-*Ci* curve) to calculate the maximum rate of ETR (J_max_). When performing these measurements in high throughput on plants acclimated to different growth environments, careful consideration needs to be given to measurement regimes to achieve accurate estimation of photosynthetic capacity (Loreto et al., 2009; Evans et al., 2017; McClain and Sharkey, 2020; Carrier et al., 1989). For instance, it is typical that HL acclimation induces a considerable reduction in the size of light-harvesting antenna linked to PSII, which, if not considered during the calculation of ETR_max_, can lead to an overestimation of the increase in photosynthetic capacity in HL-grown leaves relative to LL-grown leaves (Strand, 2014). Thus, multiple, independent methodologies for quantifying photosynthetic capacity can be used to strengthen conclusions regarding acclimation to the growth environment (Walters et al., 2003; McClain and Sharkey, 2020).

Multiple days of growth in HL are required before increased photosynthetic capacity is observed (Athanasiou et al., 2010; Walters et al., 2003). In the model herbaceous dicot *Arabidopsis thaliana*, certain genotypes can respond to a shift into HL with a leaf thickness-independent increase in photosynthetic capacity in four to five days (Athanasiou et al., 2010). That this increased photosynthetic capacity is leaf thickness-independent is important because long-term acclimation to HL for most dicots includes the development of thicker leaves (Björkman, 1981), as a consequence of additional palisade layers and/or longer palisade cells (Hanson, 1917; Evans and Poorter, 2001). Photosynthetic capacity in plants is typically expressed on a leaf area basis (m^−2^), thus the thicker leaves developed in HL have higher photosynthetic capacity than thin leaves developed in LL. Normalization of photosynthetic capacity by dry leaf weight per unit area (DW^−1^), a value that closely tracks with leaf thickness, can provide an alternative methodology to compare photosynthetic capacity between plants with different leaf thicknesses (Stewart et al., 2017; Givnish, 1988; Evans and Poorter, 2001). Species and ecotypes that exhibit strong acclimation of photosynthetic capacity use both leaf thickness-dependent and leaf thickness-independent (cell biological) mechanisms to increase photosynthetic capacity in HL, a group that includes certain ecotypes of *A. thaliana* as well as diverse species such as *Atriplex triangularis*, *Urtica dioica*, and *Radyera farragei,* whereas other species and ecotypes appear to exclusively use leaf thickness-dependent mechanisms to increase photosynthetic capacity in HL (Evans and Poorter, 2001; Murchie and Horton, 1997; Givnish, 1988; Athanasiou et al., 2010). A comparative approach between species and ecotypes with different acclimation responses to HL provides a path towards identifying the signaling pathways and metabolic pathways underlying the cell biological contributions to increased photosynthetic capacity in HL-grown plants.

The acclimation of photosynthetic capacity to HL depends on a plant’s dynamic metabolic response of foliar metabolism (Demmig-Adams et al., 2014; Häusler et al., 2014; Herrmann et al., 2019; Ma et al., 2014). In addition to carbon (C) signals, the nitrogen (N) state of the plant is also a key regulatory node controlling the acclimation of photosynthetic capacity to HL (Evans and Poorter, 2001; Paul and Pellny, 2003). Primary nitrogen assimilation is induced in HL as the enhanced PET rate increases the supply of reducing equivalents in the chloroplast (Foyer et al., 2003; Osmond et al., 1997). Nitrogen assimilation depends on the activity of chloroplastic enzymes glutamine synthetase (GS) and glutamine-oxoglutarate glutamate aminotransferase (Fd-GOGAT) (Fig. S1A). Thus, increased nitrogen assimilation rates in HL can be observed as an increase in glutamine (Gln)- and glutamate (Glu)-derived amino acids (AAs), and in particular, as an increase in the Gln/Glu ratio in HL in a manner that is proportional to the rate of PET in HL (Foyer et al., 2003, 1994). Linking C and N metabolism, the maintenance of high N assimilation rates in HL-grown plants will depend on an enhanced supply of organic acids (OAs) to the chloroplast (Nunes-Nesi et al., 2010; Dutilleul et al., 2005). These OAs can either be supplied via the *de-novo* synthesis of OAs through anaplerotic reactions or the mobilization of existing stores of OAs in leaves (Pracharoenwattana et al., 2010).

Enhanced rates of photorespiration are intertwined with increased primary nitrogen assimilation rates in HL (Rachmilevitch et al., 2004). Photorespiration rates in HL increase as a consequence of high PET and Rubisco carboxylation rates that typically drive down intracellular CO_2_ levels without a commensurate decrease in O_2_ levels (Ma et al., 2014; Keeley and Rundel, 2003). For example, in *A. thaliana,* acclimation to HL roughly doubled the carboxylation rate, whereas photorespiratory flux increased by ∼3.3 times (Ma et al., 2014). Additionally, triose phosphate utilization pathways are often rate-limiting for photosynthesis in HL, and in such circumstances, photorespiration acts as a critical photoprotective pathway and valve to convert excess sugar-phosphates into AAs (Busch et al., 2018; Wingler et al., 2000; Dirks et al., 2012). In much the same way as Gln/Glu can be used as a proxy for nitrogen assimilation and PET rates, the ratio of glycine (Gly) to serine (Ser) can be used as a metabolic indicator of photorespiration rates (Foyer et al., 2003). 2-phosphoglycolate generated by photorespiration is converted into the Gly in the peroxisome, and then in the mitochondria, glycine decarboxylase (GDC) converts Gly into Ser (Fig. S1B), which can be a rate-limiting step for photorespiration under strong photorespiratory conditions (Timm et al., 2012). The GDC reaction also releases NH_4_^+^ as a byproduct, which is reassimilated predominantly via the chloroplastic GS and Fd-GOGAT enzymes, again highlighting the interconnectedness of the N assimilation and photorespiration pathways (Leegood et al., 1995). Recently it has been shown that photorespiratory metabolic signals can repress photosynthesis, and whether this type of signal might influence the acclimation of photosynthetic capacity to HL is unknown (Li et al., 2019; Flügel et al., 2017).

In this work, the increase in photosynthetic capacity in response to a shift to HL growth is measured in *A. thaliana* using a phenotypically diverse panel of genome-sequenced natural accessions (Cao et al., 2011) and supplemented by a detailed investigation of three well-studied *A. thaliana* accessions (Ågren and Schemske, 2012). To identify metabolites linked to photosynthetic capacity acclimation, metabolomic data was generated for an ecotype with strong photosynthetic capacity acclimation to HL and a second ecotype with weak acclimation to HL. The relationship of N assimilation and photorespiration rates to the acclimation of photosynthetic capacity to HL was tested by quantifying metabolic indicators of N assimilation (Gln/Glu) and photorespiration (Gly/Ser) for the complete ecotype panel during the shift to HL. Further, ecotypes and photorespiratory mutants (Somerville and Ogren, 1980, 1981) were shifted into HL in low O_2_ and high O_2_ conditions, thereby allowing for quantification of acclimation to photosynthetic capacity acclimation in HL environments with modified rates of photorespiration. Our findings demonstrate a positive coupling between (1) indicators of N assimilation and photorespiration rate early in the HL shift, including GS activity, and (2) the magnitude of the increase in photosynthetic capacity later in the HL shift. Multiple, independent lines of evidence converge on the conclusion that low photorespiratory capacity early in the HL shift is sufficient to block the induction of photosynthetic capacity acclimation to HL.

## Methods

### Plant material and growth conditions

*Arabidopsis thaliana* ecotypes Castelnuovo-12 (IT) [ABRC stock number: CS98761], Rodasen-47 (SW) [ABRC stock number: CS98762], Col-0, and a set of 80 genome-sequenced natural accessions (Cao et al., 2011) [ABRC stock number: CS76427] were germinated following a one-week dark-cold treatment on ½ MS plates (Murashige & Skoog with Vitamins, Caisson Labs, Smithfield, UT) at 100 μmol photons m^−2^ s^−1^ in a Percival Scientific E36HID growth chamber (Perry, IA, USA) and transferred to 2.5-inch square pots with a 3.5-inch depth containing Sungro Sunshine Mix 4 (Agawam, MA, USA) 10 days post-germination. For more information on the IT and SW ecotypes, see (Oakley *et al*., 2014; Stewart *et al*., 2017). Two lines from the set of 80 accessions [ABRC stock number: CS76427] were absent from the collection at the time of this work [Mer6, ABRC stock number: CS76414 & Aitba-2, ABRC stock number: CS76347], and two lines did not have sufficient germination events to obtain an acceptable number of replicates [Castelfeld-4-212, ABRC stock number: CS76355 & Lecho-1, ABRC stock number: CS76731]; consequently, the responses of 76 of the 80 accessions were determined. Potted seedlings were grown for an additional 4 weeks in a shaded, short-day greenhouse (8-h photoperiod at 20°C/17.5°C [light/dark] with supplemental lighting (Lumigrow Pro 325, Emeryville, CA, USA) to maintain a daytime light level between 80 and 120 μmol photons m^−2^ s^−1^. Six accessions that had small rosettes under the pre-HL shift conditions were grown for 10 additional days to produce sufficient biomass before the HL shift (Bozen-1.2, Leb-3, Kly-1, Shigu-1, Koz-2, Shigu-2). Plants were randomly shuffled each day to prevent positional effects, watered every other day, and fertilized once per week using Professional 20-20-20 (Scotts Company, Marysville, OH, USA). LL-grown and HL-shifted plants were transferred in parallel for 4 days to 100 μmol photons m^−2^ s^−1^ and 800 μmol photons m^−2^ s^−1^ in a Conviron E15 growth chamber (Controlled Environments Ltd., Manitoba, Canada) with the same day-length and ambient temperatures.

### Chemicals and reagents

Standards for the metabolomic study were purchased from Sigma (St. Louis, MO, USA). [U-^13^C_4_]fumarate, [U-^13^C_2_]glycine, and [U-^13^C_6_]glucose were purchased from Isotec (Miamisburg, OH, USA). LC-MS grade acetonitrile was purchased from Thermo Fisher(Pittsburgh, PA, USA). For targeted metabolite and enzymatic assays, Amplex Red was purchased from Thermo Fisher Scientific (Waltham, MA, USA), and all enzymes were purchased from Sigma except glycine (Gly) oxidase, D-serine (Ser) dehydratase, and L-Ser racemase (BioVision, Milipitas, CA, USA).

### Leaf phenotypic traits

Photosynthetic electron transport rates were estimated by the equation ETR = ΦPSII x α x β x *I* x γ, where ΦPSII is the operating efficiency of photosystem II (PSII), α is the leaf absorptance value, β is the fraction absorbed by PSII, *I* is the photon flux density applied to the sample, and γ is the relative efficiency of light absorption of blue and red light (Loreto et al., 2009; McClain and Sharkey, 2020). The variables β and γ were assumed to be constant across our samples (0.5 and 0.75, respectively) given the identical light quality for the growth regime of all plants and shared light quality and intensity used for all samples in the measurement regime (Mekala et al., 2015). Leaf absorptance (α) was estimated using the empirically derived formula first proposed by Evans (Evans, 1996) as Chl per unit area/(Chl per unit area + 0.076), in which Chl per unit area was expressed in units of mmol m^−2^. ΦPSII was measured in tandem with q_L_, a measure of the oxidation state of the primary electron acceptor of PSII, Q_A_, using a pulse-amplitude-modulated (PAM) chlorophyll fluorometer (FMS2; Hansatech Instruments Ltd., Norfolk, UK) as previously described (Brooks and Niyogi, 2011). Briefly, light-acclimated leaves were subjected to 6.5 min of strong, white LED actinic light (1650 μmol photons m^−2^ s^−1^) and steady-state fluorescence (F_s_) levels were recorded. Maximum fluorescence levels (F_m_′) were obtained by applying a saturating pulse of light (0.8 s of 3000 µmol photons m^−2^ s^−1^) at 3 min, 5 min, and 6.5 min, and minimum fluorescence levels (F_o_′) were recorded by briefly darkening the leaf, followed by a short far-red light pulse. Q_A_ oxidation state was calculated as q_L_ = (1/F_s_ − 1/F_m_′)/(1/F_o_′ − 1/F_m_′). ΦPSII was calculated as ΦPSII = (F_m_′ - F_s_)/F_m_′. Maximum O_2_ evolution rates were determined as light- and CO_2_-saturated O_2_ evolution with leaf-disc O_2_ electrodes (Hansatech Instruments Ltd., Norfolk, UK) as previously described (Delieu and Walker, 1981). Gas exchange measurements were performed using the LI-COR 6400XT (LI-COR, Lincoln, NE) as previously described (Long and Bernacchi, 2003; Sharkey, 1988). Saturating light levels, as well as saturating CO_2_ levels, were empirically determined using the SW ecotype before commencing this work (Fig. S1). Chl *a* and *b* content per unit area was determined via spectrophotometry as previously described (Ritchie, 2006, 2008) using a methanol extraction from freeze-dried and ground leaf discs, collected after a 2-h dark period. Dry leaf mass per unit area (DW^−1^) was measured using leaf discs freeze-dried for 48 h.

### Metabolite extractions

Leaves for all metabolite extractions were flash-frozen within the growth chamber taking care to prevent shade effects at midday. Metabolites for the metabolomic study were extracted from 5 mg of freeze-dried, ground leaves using boiling water as previously described (Cocuron et al., 2014; Cocuron and Alonso, 2014). Internal standards, 100 nmol of [U-^13^C]glucose, 50 nmol of [U-^13^C]glycine, and 10 nmol of [U-^13^C]fumarate, were added at the time of the extraction. After lyophilization, extracts were resuspended in 350 µL of double distilled, deionized H_2_O, and vortexed. To quantify sugars and sugar alcohols, 150 µL of the sample was loaded onto a 0.2 µm MF centrifugal device (Nanosep, New York, NY, USA). The remaining 200 µL was transferred to a 3 kDa Amicon Ultra 0.5 mL filtering device (Millipore, Burlington, MA) for the quantification of AAs, phosphorylated compounds, and OAs. Samples were centrifuged at 14,000×g for 45 min at 4^°^C. Photorespiratory intermediates were extracted for targeted studies from 10 mg of freeze-dried, ground leaves using 200 µL H_2_O at 100°C for 5 min. After centrifugation and collection of the supernatant, this step was repeated and the two supernatants were combined for a final volume of ∼ 400 µL. Extractions for enzymatic assays were performed on 50 mg (fresh weight) ground leaves as previously described (Gibon et al., 2004).

### Quantification of intracellular metabolites and enzymatic rates

For the metabolomic work, intracellular metabolite levels (AAs, OAs, phosphorylated compounds, sugars, and sugar alcohols,) were quantified using ultrahigh-performance liquid chromatography (UHPLC) 1290 Infinity II from Agilent Technologies (Santa Clara, CA, USA) coupled with a 5500 QTRAP mass spectrometer (LC-MS/MS) from AB Sciex Instruments (Framingham, MA, USA) as previously reported (Cocuron et al., 2014) with minor modifications. Briefly, for sugars and sugar alcohols quantification, the extracts from leaves obtained after centrifugation in Nanosep MF filter were diluted 100X in acetonitrile/water (60:40, v/v) solution, then 5 µL of the diluted sample was injected onto the LC-MS/MS column. For AAs, the filtered extracts were diluted 50X in 1 mM hydrochloric acid, and 2.5 µL of the diluted sample was analyzed by LC-MS/MS. To quantify phosphorylated compounds and OAs, the filtered extracts were diluted 10X in ultra-pure water, and 10 µL was injected onto the column. LC-MS/MS data were acquired and processed using Analyst 1.6.1 software. The quantification of each intracellular metabolite was performed: i) using [U-^13^C_4_]fumarate, [U-^13^C_2_]glycine, and [U-^13^C_6_]glucose as internal standards to account for any loss of material during sample extraction and preparation; and ii) by correlating the resulting peak area of each metabolite with its corresponding external standard injected at a known concentration.

Targeted quantification of photorespiratory intermediates glutamine (Gln), glutamate (Glu), Ser, and Gly was performed by enzyme-coupled fluorescence assays with Amplex Red. Standard curves were created from 0 to 100 uM in H_2_0 for each metabolite. Gln samples in duplicate were prepared by treating 10 µL of extracted AAs and standard curve with 10 µL of 2.7 U of glutaminase (Megazyme, Bray, Ireland) in 40 mM NaAcetate, pH = 4.9 for 20 min at 37°C. On the same 96-well plate, duplicated 10 µl Glu and standard curve samples were mixed with 10 µL 40 mM NaAcetate, pH = 4.9. Fluorescence was evolved in 180 µL of 0.25 U/mL horseradish peroxidase (HRP), 27.8 mU/mL glutamate oxidase, and 0.1 mM Amplex Red in 100 mM Tris pH=7.5 for 30 min. In separate wells, paired Glu samples were treated with 180 µL containing 0.25 U/mL HRP and 0.1 mM Amplex Red in 100mM Tris pH = 7.5 for 30 min. Fluorescence was quantified (Tecan, Männedorf, Switzerland) at an excitation of 545 nm, bandwidth 10 nm, and emission detection at 590 nm, bandwidth 5 nm. Ser samples in duplicate were prepared by treating 10 µL of extracted AAs and standard curve with 10 µL of 32 µU L-SER racemase, 40 µM pyridoxal 5′-phosphate (PLP), and 50 mM NaCl in 50 mM KPO_4_, pH = 7.4 for 30 min at 37°C. Negative control samples were treated identically except L-Ser racemase was absent. Fluorescence was measured in 180 µL of 10 µU of D-Ser dehydratase, 20 µM PLP, 5 µM ZnCl_2_, 0.25 U/ml HRP and 0.1 mM Amplex Red in KPO_4_, pH = 7.4 + 50 mM NaCl for 30 min at 37°C. Gly was quantified in duplicate, along with standard curve samples, by combining 10 µL of extracted AAs with 180 µL 3.5 mU GLY oxidase, 0.25 U/ml HRP, and 0.1 mM Amplex Red in 20 mM KPO_4_, pH = 9, 100 mM NaCl, and 10% glycerol (v/v) for 30 min at 37°C. Negative controls were treated identically except Gly oxidase was absent.

Enzymatic rates for Gln synthetase (GS), ferredoxin-glutamine-oxoglutarate glutamine transferase (Fd-GOGAT), PEP carboxylase, pyruvate kinase, and cytosolic fructose bisphosphatase were measured as previously described (Gibon et al., 2004) with the minor modification that 20 µL of the extract was used and specifically for GS, the catalytic rate was quantified after stabilizing for 5 min.

## Results

### Variation in acclimation of photosynthetic capacity to HL among natural accessions

LL-grown plants of the reference *A. thaliana* genotype Col-0 and two ecotypes locally adapted to contrasting environments (IT and SW) were shifted into HL, and photosynthetic capacity measurements were taken over 10 days along with measurements of the Q_A_ redox state, as quantified by the chlorophyll fluorescence parameter q_L_. A statistically significant increase in q_L_, when measured in saturating light, preceded the increase in photosynthetic capacity (q_L_ on Day 3 vs. ETR_max_ and P_max_ on Day 4) (Fig. 1A). The fastest rate of increase in q_L_ occurred between HL Day 1 and Day 3, whereas photosynthetic capacity on a leaf area basis (m^−2^) rose most sharply between HL Day 4 and Day 6. In contrast, the increase in photosynthetic capacity plateaued at Day 4 when normalized by DW^−1^, a proxy for leaf thickness. This indicates that by HL Day 6, leaves that had developed in HL showed the expected increase in leaf thickness (Stewart et al., 2017). The fastest rate of increase in P_max_ normalized by DW^−1^ (i.e., leaf thickness-independent increases in photosynthetic capacity) occurred between Day 3 and Day 4 of the post-HL shift (Fig. 1A).

**Figure 1.**
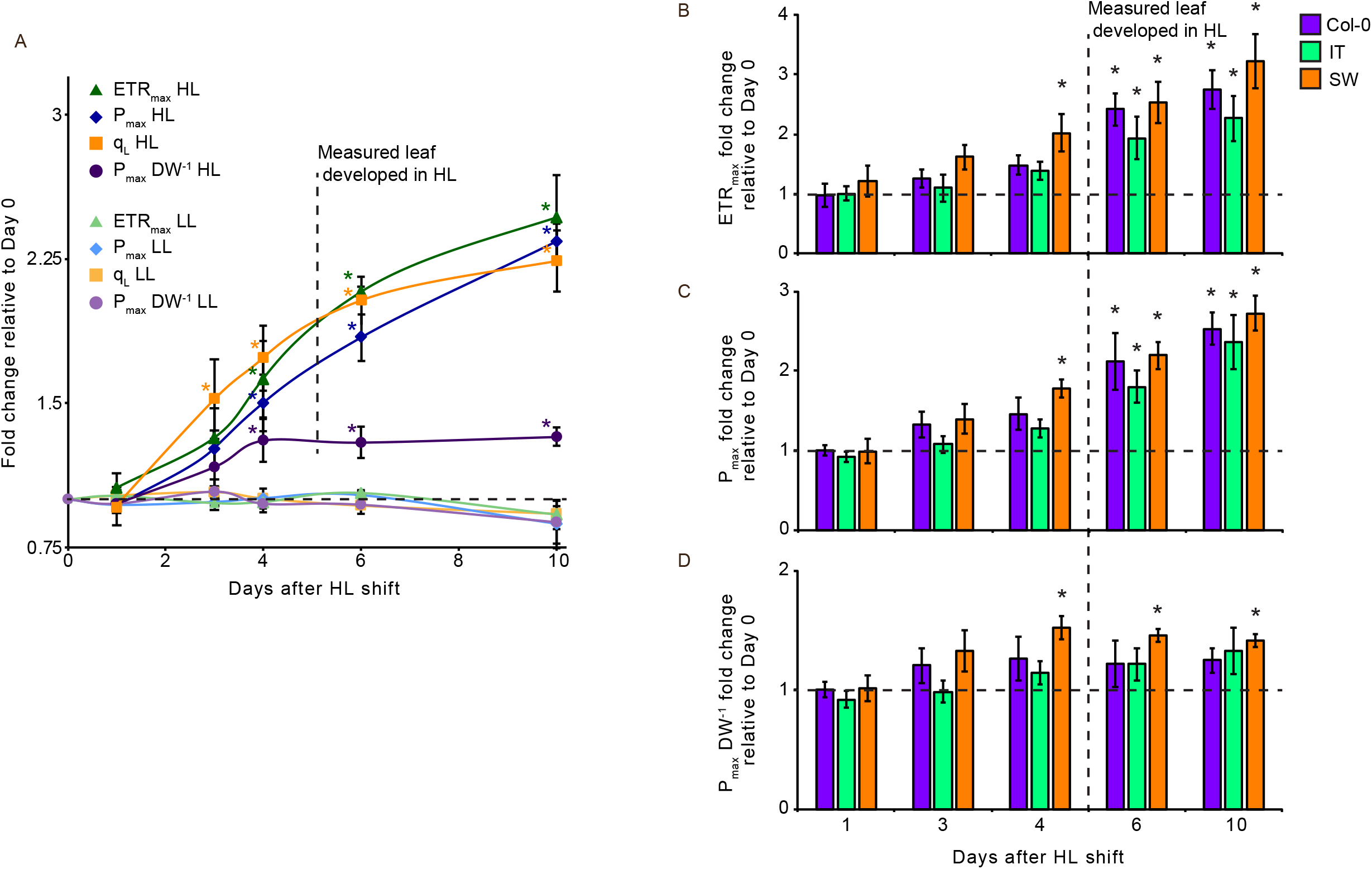
The dynamics of photosynthetic capacity acclimation to HL in the Col-0, IT, and SW plants. (a) Photosynthetic capacity as measured by maximum electron transport rate, ETR_max_ (i.e., maximal light-saturated photosynthetic electron transport rate) per leaf area (green), maximum oxygen evolution rate, P_max_ (i.e., maximal CO_2_- and light-saturated oxygen evolution rate) rate per area (blue), and P_max_ per unit dry leaf mass (DW^−1^) (purple). Q_A_ redox state was measured as q_L_ (orange). Mean values combine data from Col-0, IT, and SW plants ± standard errors (*n* = 12). (b) ETR_max_ per unit area, (c) P_max_ per unit area, and (d) P_max_ per DW^−1^ for the Col-0, IT, and SW plotted individually. Mean values ± standard errors (*n* = 4) are given an asterisk if the mean value at the indicated time point was significantly different from the mean value for that accession at Day 0 based on a one-way ANOVA and post hoc Tukey–Kramer HSD tests.

While the three accessions shared a similar response of q_L_ and photosynthetic capacity to the shift to HL growth, the SW ecotype acclimated to HL growth condition faster than the other two accessions (Fig. 1B-D, S2). In particular, the SW ecotype on HL Days 3 and 4 had higher photosynthetic capacity relative to the other two accessions. Post-HL-shifted SW ecotype plants also achieved the highest P_max_ normalized by DW (Fig. S2D). The more rapid acclimation of the SW ecotype was also observed when q_L_ was measured using the growth light intensity (Fig. S2E). In contrast to q_L_ measured at saturating light, q_L_ at the growth light intensity significantly increased in both the Col-0 and SW on HL Day 1, with the larger magnitude increase in the SW ecotype (Fig. S2E). Thus, we conclude that the SW ecotype possesses a more robust initial phase of the photosynthetic capacity acclimation response relative to the Col-0 and IT accessions.

To map the natural diversity of the photosynthetic acclimation response to HL in greater detail, HL shifts were performed on LL-grown plants of the geographically and phenotypically diverse genome-sequenced *A. thaliana* ecotype population first assembled by Cao et al. (Cao et al., 2011). HL Day 4 was selected as an optimal point to test ecotypic variation in the acclimation of photosynthetic capacity to HL due both to the strong acclimation of photosynthetic capacity in the SW ecotype relative to Col-0 and IT and the absence of changes in DW in those three accessions at that time point (Fig. 1, S2). A range of fold changes in photosynthetic capacity (HL Day 4/LL) was observed from a low of ∼0.95 to a high of ∼2.2-fold (Fig. 2). Accessions that showed strongly increased photosynthetic capacity, such as Dog-4 (ecotype #5), had a fold change in photosynthetic capacity that exceeded the fold change of the SW ecotype when quantified by either ETR_max_ or P_max_. At the opposite extreme, some ecotypes showed virtually no change or even a slight decline in HL when the fold change in photosynthetic capacity (HL Day 4/LL) was measured by either ETR_max_ or P_max_, best exemplified by the ecotypes Bak-2 and Lago-1 (ecotypes #70 and #78, respectively).

**Figure 2.**
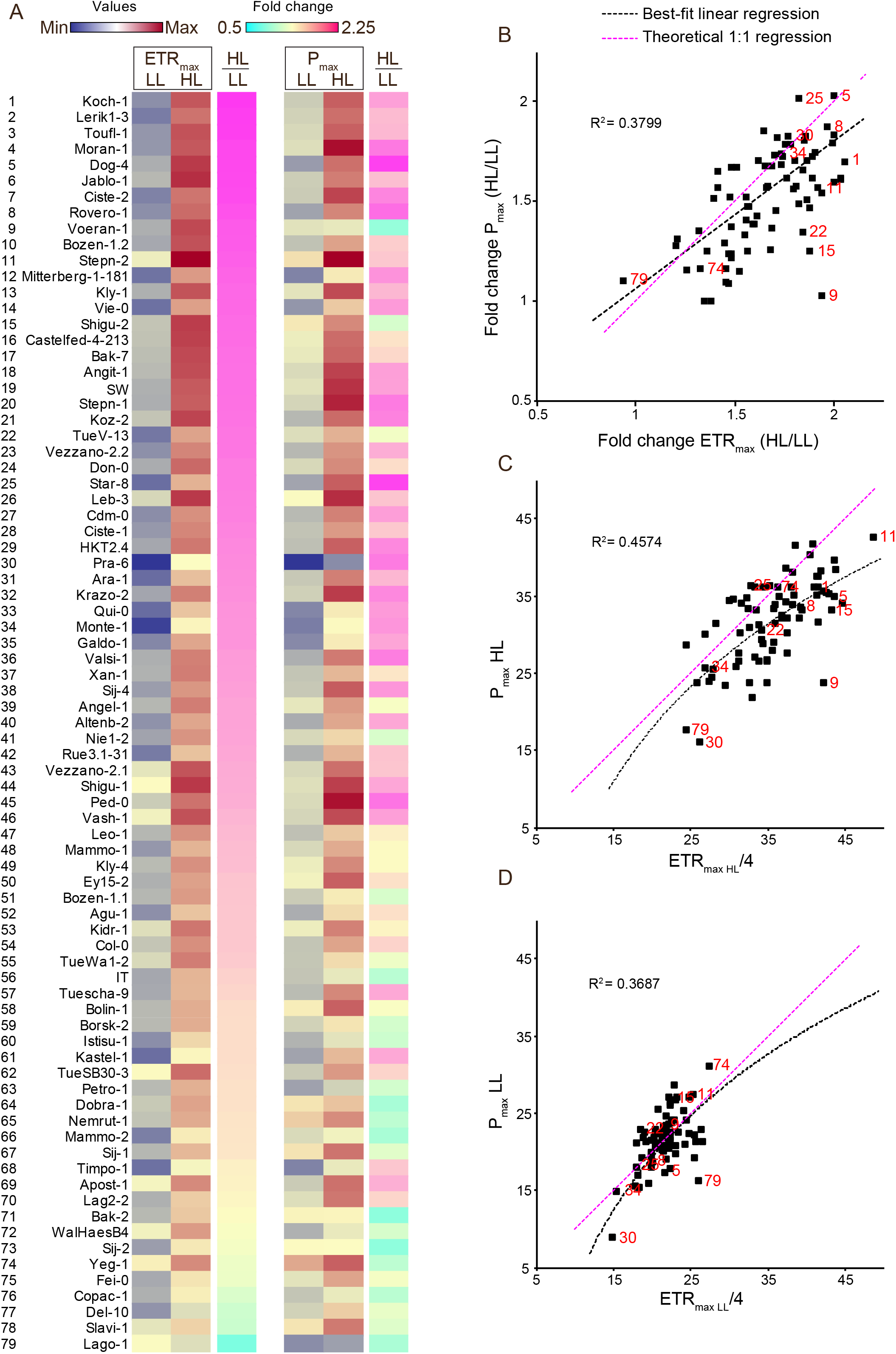
Natural diversity in photosynthetic capacity acclimation response to HL. (a) Photosynthetic capacity as measured by ETR_max_ and P_max_ per leaf area for Col-0, IT, SW, and the accession panel (Cao et al., 2011) plants from LL growth (LL) and on HL Day 4 (HL). LL and HL plants are displayed on a shared color-scale for ETR_max_ and P_max_, respectively, from low (dark blue) to high (carmine red). Fold-change (HL/LL) values are shown on a separate color scale with a range of 0.5 (cyan) to 2.25 (magenta). Each ecotype is assigned a unique number in descending order of the magnitude of the HL/LL ETR_max_ fold-change. (b-d) Correlation of P_max_ with ETR_max_/4 data for (b) HL/LL fold-change, (c) HL plants, and (d) LL plants. Best-fit regression and associated R^2^-values (black) and theoretical 1:1 regression (magenta) are displayed. Ecotype numbers corresponding to the unique number for each ecotype in panel a (red) are included for the outlier ecotypes. ETR_max_ and P_max_ mean values were calculated using *n* = 2 to 4 biological replicates.

Chl per unit area on average decreased to ∼75.7% in HL-shifted plants relative to LL with extremes of slight induction in Chl per unit area for ecotypes such as Shigu-1 and Stephn-2 (ecotypes #43 and #11, respectively) and a greater than 2-fold reduction in chlorophyll per unit area for ecotypes such as Don-0 and Leb-3 (ecotypes #23 and #25, respectively) (Fig. S3). Importantly, no correlation was observed between chlorophyll per unit area and ETR_max_ values (Fig. S3). This verifies that the use of chlorophyll per unit area values in HL Day 4 plants and LL as a means to estimate light absorptance for use in the calculation of the fold change in ETR_max_ did not systematically bias these calculations (see Materials & Methods for details on the estimation of ETR_max_ values).

The fold change in photosynthetic capacity (HL Day 4/LL) quantified by P_max_ and ETR_max_ across the population was tightly correlated (Fig. 2B, R^2^ = 0.3799). Similarly, the fold change in q_L_ (measured in saturating light) was also correlated with the fold change in ETR_max_ (Fig. S4, R^2^ = 0.2062), consistent with the increase in photosynthetic capacity contributing to the restoration redox poise of the PET chain in HL-shifted plants. The correlation was weaker between ETR_max_ and P_max_ or q_L_ for LL-grown plants relative to these same comparisons made in HL-shifted plants (Fig. 2C-D, Fig. S4). The average increase in photosynthetic capacity, calculated from all the accessions, as measured by ETR_max_ (ETR_max HL_/ETR_max LL_ ≈ 1.66) was higher than the average fold change of photosynthetic capacity as quantified by P_max_ (P_max HL_/P_max LL_ ≈ 1.48). Precise measurement of ETR_max_ necessitates direct quantification of values that are only indirectly estimated in this work due to the scale of phenotyping(McClain and Sharkey, 2020; Loreto et al., 2009; Evans et al., 2017; Evans, 1996). Hence, in some cases the ETR_max_ reported here may overestimate the fold change in photosynthetic capacity due potentially to changes in, for instance, PSII-to-PSI state transitions, or carotenoid contributions to light absorptance. The alternative, and mutually non-exclusive possibility, is that P_max_ can underestimate PET rates due to oxygen-consuming pathways, while significantly repressed, persisting in saturating CO_2_ ^(^Carrier et al., 1989^)^.

In this context, it is worth noting several outlier ecotypes that had ETR_max_ values that strongly contrasted with P_max_ values under one of the growth conditions. For instance, ecotypes such as Voeran-1 (ecotype #9), Shigu-2 (ecotype #15), and TueV-13 (ecotype #21) strongly increased photosynthetic capacity when quantified by ETR_max_ but had a moderate or no increase in P_max_ following the HL shift (Fig. 2B-D). Specifically, this was a consequence of low P_max_ values relative to ETR_max_ when measured in the HL-shifted plants for these three ecotypes. Whereas the majority of ecotypes showed quite similar P_max_ and ETR_max_ values, it would make an interesting case study to investigate what specific mechanisms underlie the growth condition-specific divergence of ETR_max_, as estimated in this work, and P_max_ values in ecotype such as Vorean-1.

### Impact of the HL shift on the leaf metabolome

To identify the metabolic changes linked to the increase in photosynthetic capacity during the HL shift, metabolomic data were collected for the SW ecotype, a strongly acclimating ecotype, and Col-0, a weakly acclimating ecotype. A principal component (PC) analysis of HL-shifted and LL-grown plants for both ecotypes separated both Col-0 from SW plants and HL-shifted from LL-grown plants on the principal component 1 (PC1) axis (Fig. S5A). A PC analysis of just SW plants separated HL-shifted from LL-grown plants along PC1 and then separated along the time-axis of the HL treatment on PC2 (Fig. S5B). Thus, the metabolomes could be reliably resolved by ecotype, growth condition, and the time component.

Broadly, sugar-phosphates, sugars, and AAs had increased abundance in HL-shifted plants (Fig. 3, best illustrated by the left set of columns). The exceptions to this overall trend of increased abundance in HL-shifted plants included the decreased abundance of the AAs lysine (Lys), the photorespiratory intermediate Ser, threonine (Thr), as well as in the SW ecotype the glycolytic intermediate phosphoenolpyruvate (PEP) (Fig. 3, best illustrated by the left set of columns). AAs, as a class of metabolites, increased over time in response to the HL shift driven primarily by the increase over time of the high abundance Glu- and Gln-derived AAs (Fig. 3, best illustrated by the left set of columns). The high abundance foliar metabolites to have the inverse profile, decreasing abundance over time in response to the HL shift, were intermediates of the CBB cycle and sucrose biosynthesis (fructose 1,6-bisphosphate (FBP), sucrose 6-phosphate (Suc6P), and ribulose 1,5-bisphosphate (RuBP)), the TCA cycle (fumarate, isocitrate, and α-ketoglutarate), as well as the AAs Thr and Ser (Fig. 3, best illustrated by the middle set of columns). In contrast to the temporal response of AAs, soluble sugars reached their highest abundance on HL Day 1 driven by a transient increase in Glc and Fru levels, whereas Suc levels remained at an elevated abundance throughout the post-HL shift period (Fig. 3, best illustrated by the middle set of columns). These trends were mirrored by enzyme activity measurements in HL shifted plants for PEP carboxylase, pyruvate kinase, GS, and cytosolic fructose bisphosphatase (Fig. S6). PEP carboxylase activity was induced on HL Day 1, which is likely critical to achieve the enhanced OA synthesis rates necessary for increased AA synthesis in HL growth, while simultaneously cytosolic fructose bisphosphatase activity was reduced in concert with the increased abundance of FBP and other sucrose biosynthesis intermediates (Fig S6A,D).

**Figure 3.**
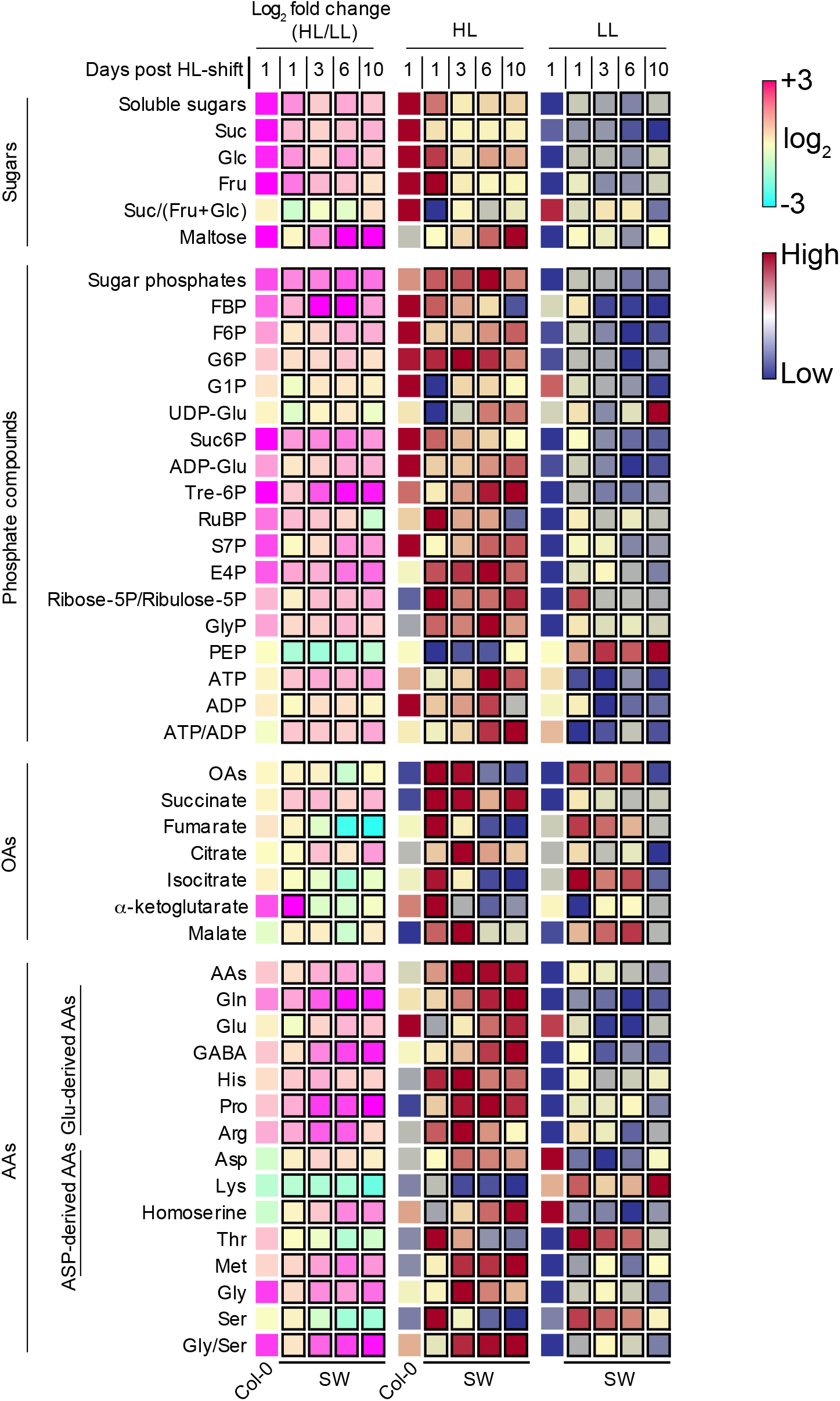
Leaf metabolomic data for metabolite classes and selected individual metabolites. Abundance of total soluble sugars, sugar phosphates, organic acids, and amino acids (AAs) classes displayed alongside selected metabolites (metabolites with an HL/LL abundance change) from each class for Col-0 (no outside square) and SW (black squares). The abundance of the metabolite for LL-grown and HL-shifted plants are displayed on a shared color-scale, low (dark blue) to high (carmine red). Log_2_-transformed HL/LL fold changes are displayed on a separate color scale from −3 (cyan) to +3 (magenta). Mean values are calculated from *n* = 3 to 8 biological replicates.

Variable importance (VIP) analyses of the supervised partial least-squares discriminant analysis (PLSDA) were performed to identify metabolites that differentiate the HL metabolomes of these ecotypes relative to their paired LL metabolome. In the SW ecotype, VIP analysis identified Gln, both a photorespiratory intermediate and product of N assimilation, as the strongest metabolomic differentiator of HL-shifted plants from LL-grown plants (Fig. 4A, Fig. S5C). Similarly, both Gln and Gly, another key photorespiratory intermediate, were among the highest VIP scores between HL-shifted and LL-grown plants in the Col-0 ecotype (Fig. 4B, Fig. S5D). When the results of the SW VIP analysis were compared to the results of the Col-0 analysis, the higher scoring metabolites in the SW VIP analysis relative to the Col-0 analysis included ATP, valine (Val), histidine (His), Glu-derived AAs γ-aminobutyric acid (GABA) and proline (Pro) as well as phosphoenolpyruvate (PEP), which was one of few components of central metabolism with decreased abundance in the HL-shifted SW plants. Likewise, the metabolites which scored more strongly in the post-HL vs. LL VIP analysis of Col-0 relative to the SW analysis included Suc6P, ADP-glucose, RuBP, sedoheptulose 7-phosphate (S7P), the AAs phenylalanine (Phe), tryptophan (Trp), aspartate (Asp), and Gly, as well as the glucose (Glc) and fructose (Fru). Increased abundance of metabolites with high VIP scores specifically in the strongly acclimating SW ecotype such as ATP and Glu-derived residues, as well as decreased PEP, represent putative metabolic indicators of efficient photosynthetic capacity to HL. Likewise, high VIP scores specifically in the weakly acclimating Col-0 ecotype, such as increased abundance of Suc6P and ADP-glucose, may represent indicators of inefficient photosynthetic capacity acclimation to HL.

**Figure 4.**
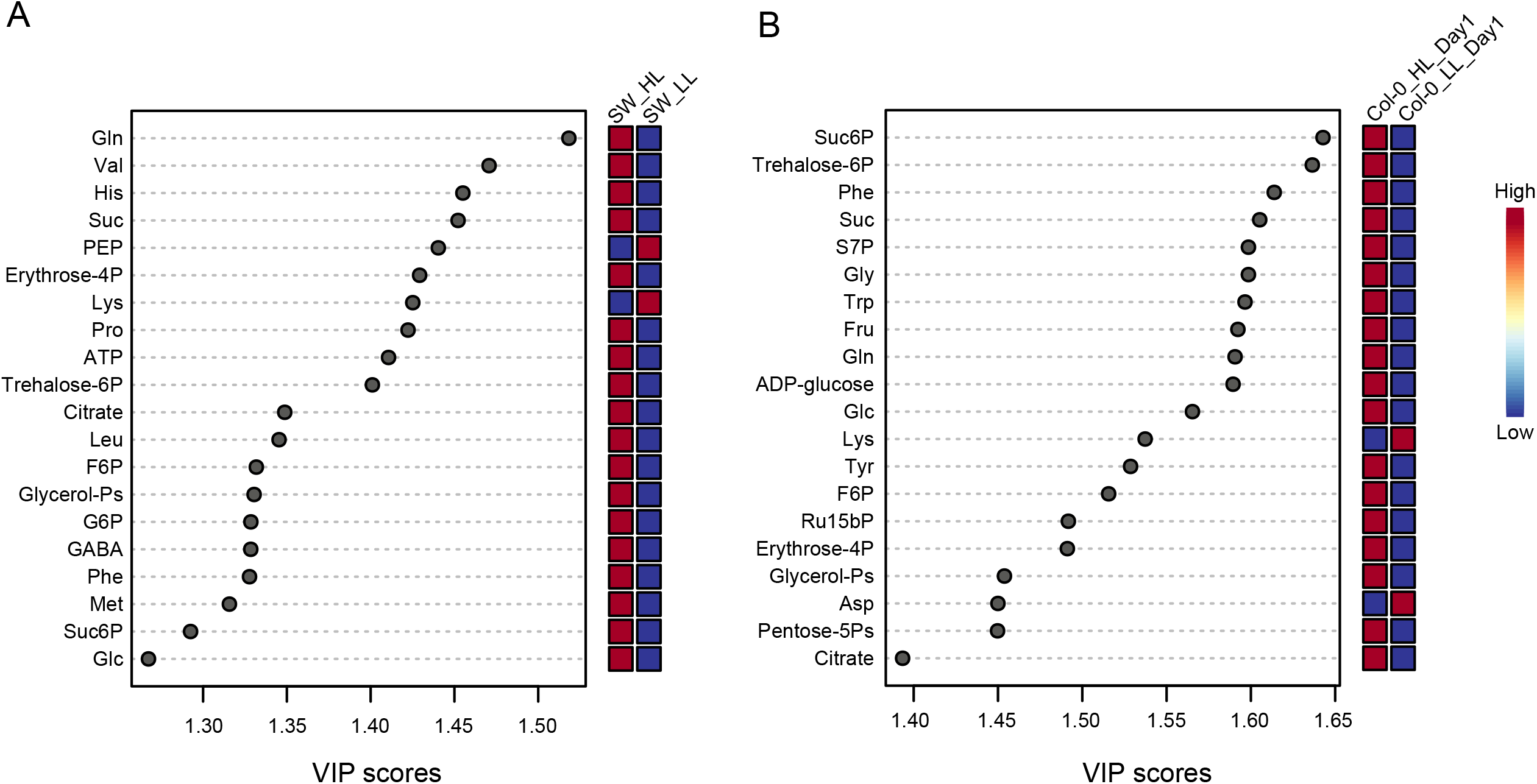
Metabolites most strongly linked to HL acclimation in the SW and Col-0 plants. Variable importance in projection (VIP) scores showing the highest 20 metabolites that contribute the most to the separation for (a) SW and (b) Col-0 plants. Relative metabolite abundance in LL and HL plants are displayed on a color scale; high abundance (carmine red) and low abundance (dark blue).

As an additional means of linking specific metabolites to the change in photosynthetic capacity, the abundance over time of metabolites in the SW ecotype was correlated with the change in photosynthetic capacity (Fig. 5). Levels of Glu, Gln, and other Glu-derived AAs such as GABA and Pro were positively correlated with photosynthetic capacity. Additionally, the Asp-derived AAs homoserine and methionine (Met) were positively correlated with photosynthetic capacity whereas the Asp-derived AA Thr was inversely correlated. The two strongest inverse correlations were for fumarate and Ser. The correlation with photosynthetic capacity of Glu- and Gln-derived AAs, products of N assimilation, highlights the relationship of this pathway with the acclimation of photosynthetic capacity to HL. The decreased abundance of metabolites involved in the supply of OAs to support N assimilation (e.g., PEP, fumarate, isocitrate, α-ketoglutarate) also fits with the importance of N assimilation rates to increased photosynthetic capacity (Fig. S1A). The strong inverse correlation of photosynthetic capacity with Ser levels and elevated Gly abundance in HL-shifted plants also indicate elevated photorespiration rates (Fig. S1B). The links between photorespiration, N assimilation, and the acclimation of photosynthetic capacity to HL were explored further in subsequent experiments.

**Figure 5.**
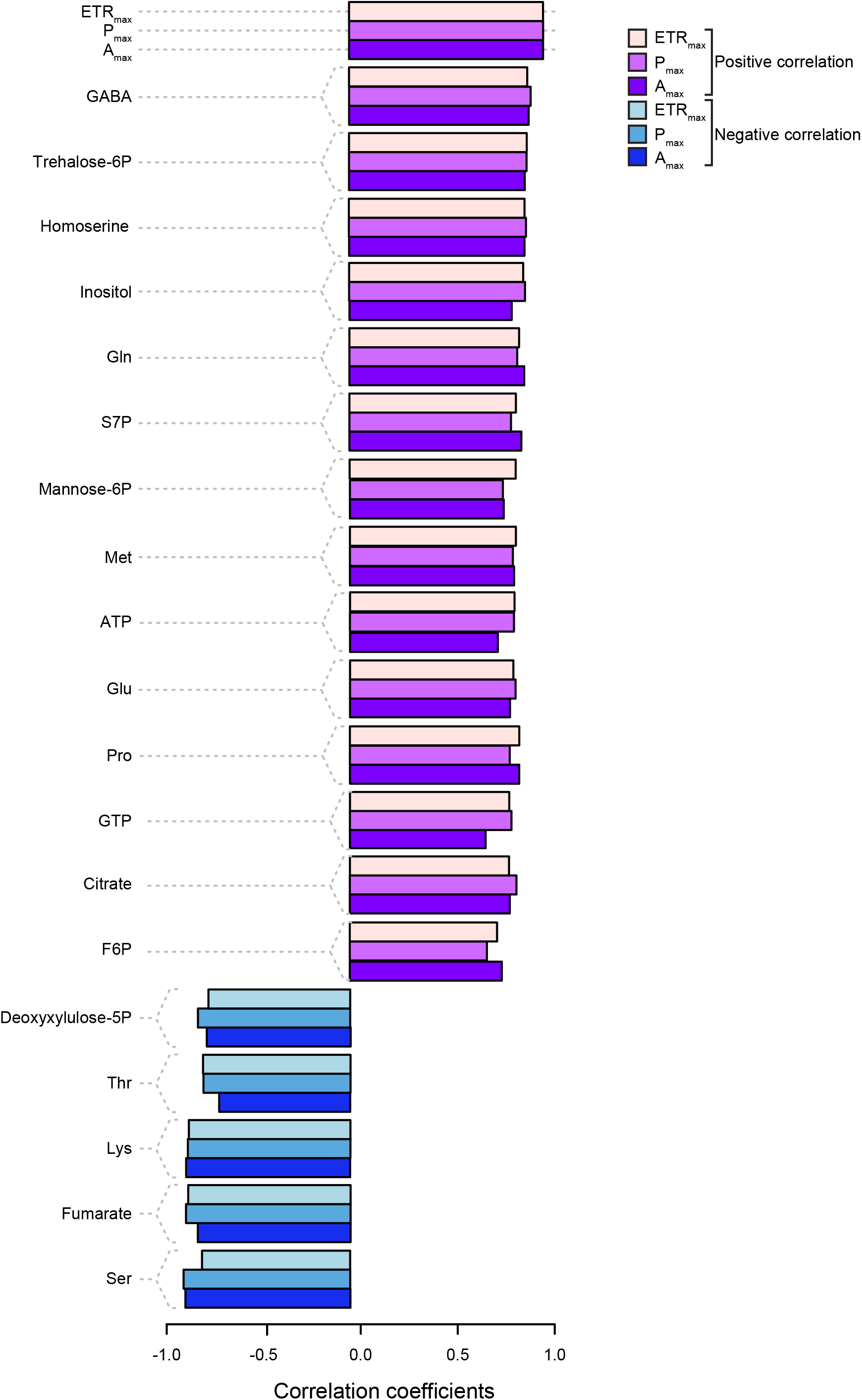
Correlation of change in metabolite abundance with photosynthetic capacity. Correlation coefficients of time-resolved metabolomics data in HL-shifted SW plants with the change in ETR_max_ per unit area, P_max_ per unit area, and A_max_ per unit area at the same sampling points; positive correlations (red) and negative correlations (blue).

### Metabolic signatures of primary N assimilation and photorespiration in HL

The SW ecotype also differed from the Col-0 and IT ecotypes in two physiological parameters relevant to photorespiration rates during the HL shift, stomatal conductance (*g_s_*) and the post-illumination CO_2_ burst (PIB) (Fig. S7). Lower *g_s_* in HL will reduce intracellular CO_2_ levels, promoting higher rates of photorespiration. *g_s_* decreased in all three ecotypes on HL Day 1, but on HL Day 3, the SW ecotype was the sole ecotype to restore pre-shift *g_s_* levels (Fig. S7A). However, the percentage increase in the net CO_2_ assimilation rate by measuring in low O_2_ air, used to repress photorespiration during gas exchange analysis, relative to ambient O_2_ air did not statistically differ between the ecotypes (Fig. S7B), indicating that the impact of *g_s_* on Rubisco oxygenation rates was not drastic. Downstream of Rubisco catalyzed RuBP-oxygenation, the magnitude of the PIB, which estimates mitochondrial GDC enzymatic capacity, increased in all three ecotypes on Day 10 of the HL shift but was increased solely in the SW ecotype on Day 3 (Fig. S7C). This supports that SW may have had higher photorespiratory capacity early in the HL shift relative to the other two ecotypes.

Whether photorespiration and primary N assimilation caused the increase in Gly and Gln levels, respectively, in HL-shifted plants was investigated by decoupling rates of these processes from the shift to HL growth conditions by combining this treatment with introducing low O_2_ air into the growth chamber. An increased Gln/Glu was observed in SW, Col-0, and IT plants on HL Day 1 in ambient O_2_ conditions with the highest Gln/Glu ratio in the SW ecotype (Fig. 6A). In contrast, at HL Day 3 in low O_2_ air, the Gln/Glu ratio decreased in all three ecotypes relative to the LL value. The total pool of Gln + Glu also increased in HL-shifted plants in ambient air and again, this increase was not observed in low O_2_ air (Fig. 6B). The Gly/Ser ratio, as well as the total pool of Gly + Ser, were also increased after the HL shift in ambient O_2_ levels in all three ecotypes and not in low O_2_ air (Fig. 6C-D).

**Figure 6.**
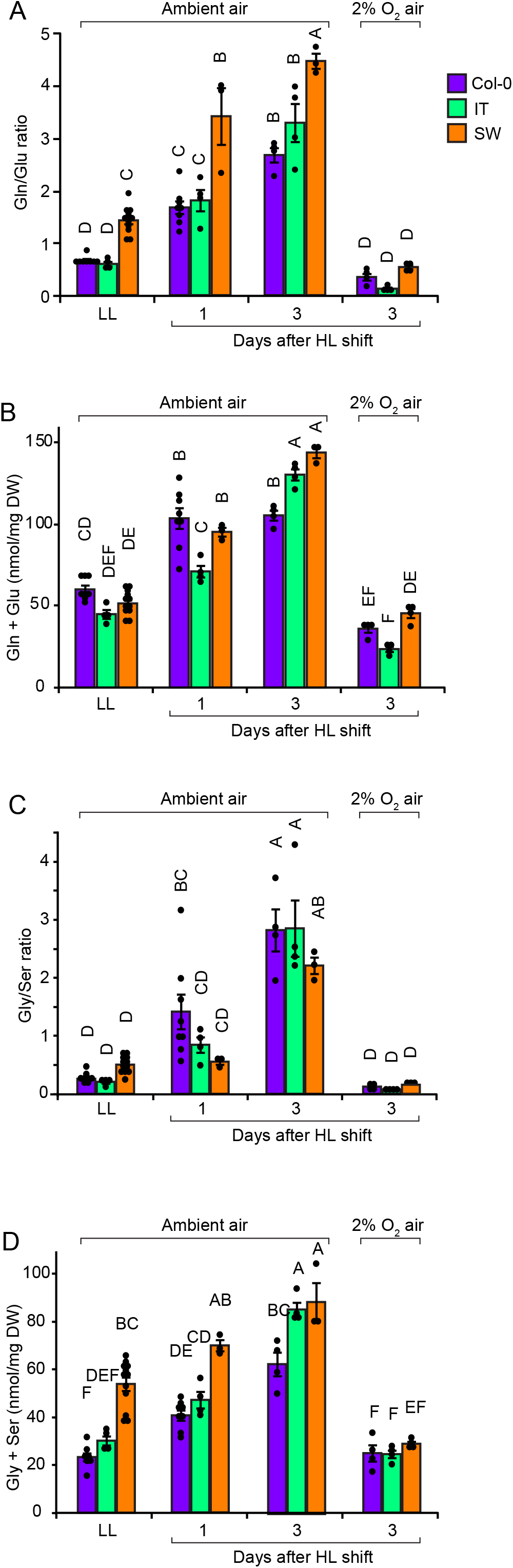
Low oxygen suppresses the HL-dependent increase in Gln/Glu and Gly/Ser. Metabolite quantification of the (a) Gln/Glu ratio, (b) nmol Gln + Glu per mg dry leaf weight (DW), (c) Gly/Ser ratio, and (d) nmol Gly + Ser per mg DW^−1^ for Col-0 (purple), IT (green), and SW (orange) plants are grown in LL and HL Day 1 and Day 3 in ambient air or Day 3 post-HL shift in low oxygen air (400 ppm CO_2_, 2% O_2_, balance N_2_). Mean values ± standard errors (*n* = 3 to 8) with individual data points displayed as dots (black). Mean values that share the same letters are not statistically different, and those that do not share the same letters are statistically different based on one-way ANOVA and post hoc Tukey–Kramer HSD tests.

While these trends are shared between the ecotypes, the Gly/Ser and Gln/Glu ratios did not behave identically in all three ecotypes. For example, the Gln/Glu ratio in ambient O_2_ conditions was consistently higher in the SW ecotype, whereas on Day 1 of the HL shift, the Gly/Ser ratio was lowest in the SW ecotype. Thus, the HL-driven increase in Gln/Glu and Gly/Ser, metabolic markers for the rates of N assimilation and photorespiration, respectively, was dependent on ambient O_2_ levels and significant differences between ecotypes for these metabolic markers could be measured on HL Day 1.

### Correlating metabolic signatures of N metabolism with the induction of high photosynthetic capacity in HL

As metabolic indicators of N assimilation and photorespiration rates, levels of Gln, Glu, Ser, and Gly, as well as the activity of enzymes GS and Fd-GOGAT (key to both primary N assimilation and photorespiration), were quantified on HL Day 1 using the full ecotype panel. The goal was to correlate changes in these metabolites on HL Day 1 with the fold change in photosynthetic capacity measured on HL Day 4. Strongly supporting a link between photorespiration and N assimilation rates early in the HL shift and the later increase in photosynthetic capacity, both the Gln/Glu and Gly/Ser ratios on HL Day 1 were positively and strongly correlated with the fold change in photosynthetic capacity on HL Day 4 (HL Day 4/LL) when measured as ETR_max_ and correlated, albeit not as strongly, with P_max_ (Fig. 7A-B, Table 1, Fig. S8, R^2^ = 0.5292 and 0.1193, respectively, for Gln/Glu, and R^2^ = 0.4034 and 0.0693, respectively, for Gly/Ser). The positive correlation between the fold change in photosynthetic capacity and the Gly/Ser ratio in the larger ecotype panel contrasted with the behavior of the smaller ecotype set of SW, Col-0, and IT plants (Fig. 6C). Additional correlations were found in this dataset, albeit none as strong as the correlation for ETR_max_ and Gln/Glu and Gly/Ser values. ETR_max_ values in HL-shifted plants were positively correlated with both the Gln/Glu and Gly/Ser ratios (Table 1, Fig. S9A-B, R^2^ = 0.1408 and 0.1670, respectively). The total Gly + Ser pool in HL-shifted plants was correlated with the fold change in photosynthetic capacity as measured by ETR_max_, as well as weakly negatively correlated with ETR_max_ for LL-grown plants (Fig. 7C, Fig. S9C, R^2^ = 0.1562 and 0.0559, respectively), again supporting a link between photorespiratory capacity and the later increase in photosynthetic capacity. In contrast to the Gly + Ser result, the total pool of Gln + Glu was not correlated with the induction of photosynthetic capacity (Table 1). GS enzymatic activity on HL Day 1 was also weakly correlated with the fold change in the photosynthetic capacity as measured by ETR_max_ and was negatively correlated with ETR_max_ for LL-grown plants (Fig. 7C, Fig. S9D, R^2^ = 0.1190 and 0.1171, respectively). GS activity is necessary for both photorespiration and primary N assimilation, thus plants with high GS activity may be indicative of either or both high photorespiratory and primary N assimilatory capacity. Similarly, GS activity on HL Day 1 was correlated with the fold change in q_L_ and was weakly negatively correlated with q_L_ for LL-grown plants (Fig. S10, R^2^ = 0.1254 and 0.0526, respectively). In summary, metabolic indicators of N assimilation, N assimilation/photorespiration enzyme activities, and metabolic indicators of photorespiration rates early in the HL shift were positively correlated with the magnitude of the subsequent increase in photosynthetic capacity measured at HL day 4

**Table 1:** Results of regression analysis for parameters in Figures 7, S7, and S8.

| Parameter | *Fold Change*<br>*(HL/LL)* |  | *HL* |  | *LL* |  |
| --- | --- | --- | --- | --- | --- | --- |
|  | ETR<sub>max</sub> | P<sub>max</sub> | ETR<sub>max</sub> | P<sub>max</sub> | ETR<sub>max</sub> | P<sub>max</sub> |
| **Gln/Glu** | *** | ** | ** | *n.s.* | **τ | *τ |
| **Gln + Glu** | *n.s.* | *n.s.* | *n.s.* | *n.s.* | *n.s.* | *n.s.* |
| *(nmol mg DW<sup>-1</sup>)* |  |  |  |  |  |  |
| **Gly/Ser** | ** | * | ** | *n.s.* | *τ | *n.s.* |
| **Gly + Ser** | ** | *n.s.* | *n.s.* | *n.s.* | *τ | *n.s.* |
| *(nmol mg DW<sup>-1</sup>)* |  |  |  |  |  |  |
| **GS** | ** | *n.s.* | *n.s.* | *n.s.* | **τ | *n.s.* |
| *(mmol ADP min<sup>-1</sup> mg protein<sup>-1</sup>)* |  |  |  |  |  |  |
| **Fd-GOGAT** | *n.s.* | * | *n.s.* | *n.s.* | *n.s.* | *n.s.* |
| *(mmol glutamate min<sup>-1</sup> mg protein<sup>-1</sup>)* |  |  |  |  |  |  |

Significant correlations are denoted by asterisks; \* = $R^2 > 0.05$ , \*\* = $R^2 > 0.1$ , \*\*\* = $R^2 > 0.5$ (*n.s.* = not significant). The symbol τ denotes an inverse correlation.

**Figure 7.**
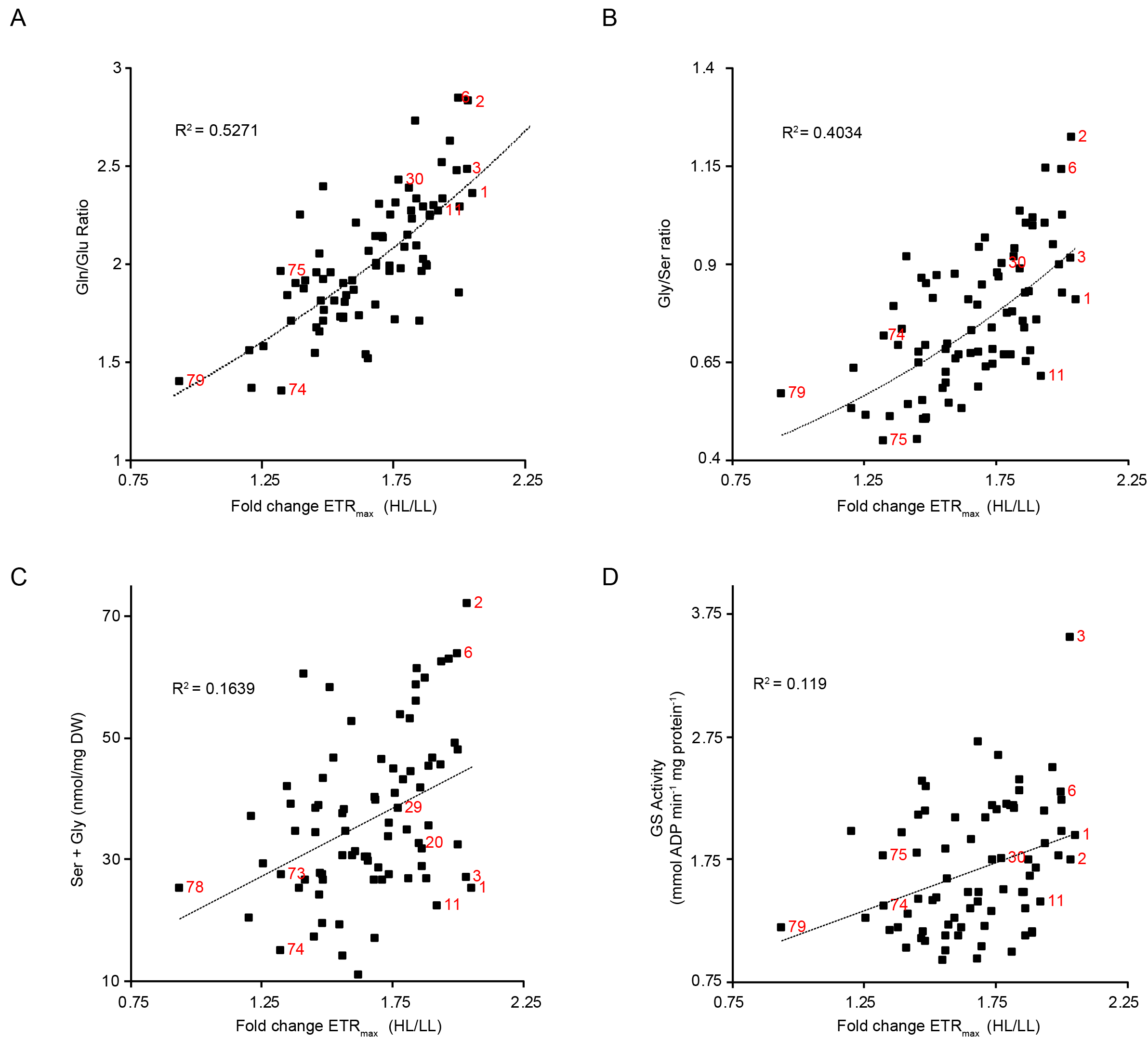
Correlation of metabolic indicators of photorespiration and nitrogen assimilation rates on HL Day 1 with increased photosynthetic capacity on HL Day 4. Fold-change (HL/LL) of ETR_max_ per leaf area for Col-0, IT, SW, and the accession panel (Cao et al., 2011) with quantification of (a) Gln/Glu, (b) Gly/ser, (c) nmol Ser + Gly per mg DW^−1^, and (d) glutamine synthetase (GS) activity (mmol adenosine diphosphate (ADP) min^−1^ mg protein^−1^) levels on HL Day 1. ETR_max_ mean values were calculated using *n* = 2 to 4 biological replicates. Metabolite and enzymatic rate quantifications were calculated using *n* = 2 to 4 biological replicates and *n* = 2 technical replicates for each biological replicate. Ecotype numbers (red) correspond to numbers in Fig. 2A.

### Manipulating photorespiration and N metabolism using O_2_ levels and impact on the induction of high photosynthetic capacity in HL

Two models can be envisioned for the relationship between increased photosynthetic capacity and photorespiration/N assimilation rates. First, high rates of N assimilation and photorespiration might generate a positive signal for photosynthetic capacity acclimation. This model predicts that a HL shift in low O_2_ air (to block photorespiration) should prevent the increase in photosynthetic capacity. Alternatively, low photorespiratory capacity could generate a negative signal that represses photosynthetic capacity acclimation to HL. This model predicts that a HL shift in high O_2_ air (or in a photorespiratory mutant) should block the increase in photosynthetic capacity. To distinguish between these models, accessions selected to cover the full range of diversity in fold change of photosynthetic capacity were shifted into HL combined with ambient air, low O_2_ air, or high O_2_ air (Fig. 8). Two mutants affecting photorespiratory enzymes, Ser hydroxymethyltransferase (*shm1*, AT4G37930) and Fd-GOGAT (*gls1-30*, AT5G04140) (Somerville and Ogren, 1981, 1980; Coschigano et al., 1998; Somerville and Ogren, 1982), were also included in this experiment to test the response of photosynthetic capacity to HL in genotypes with extremely low photorespiratory capacity.

**Figure 8.**
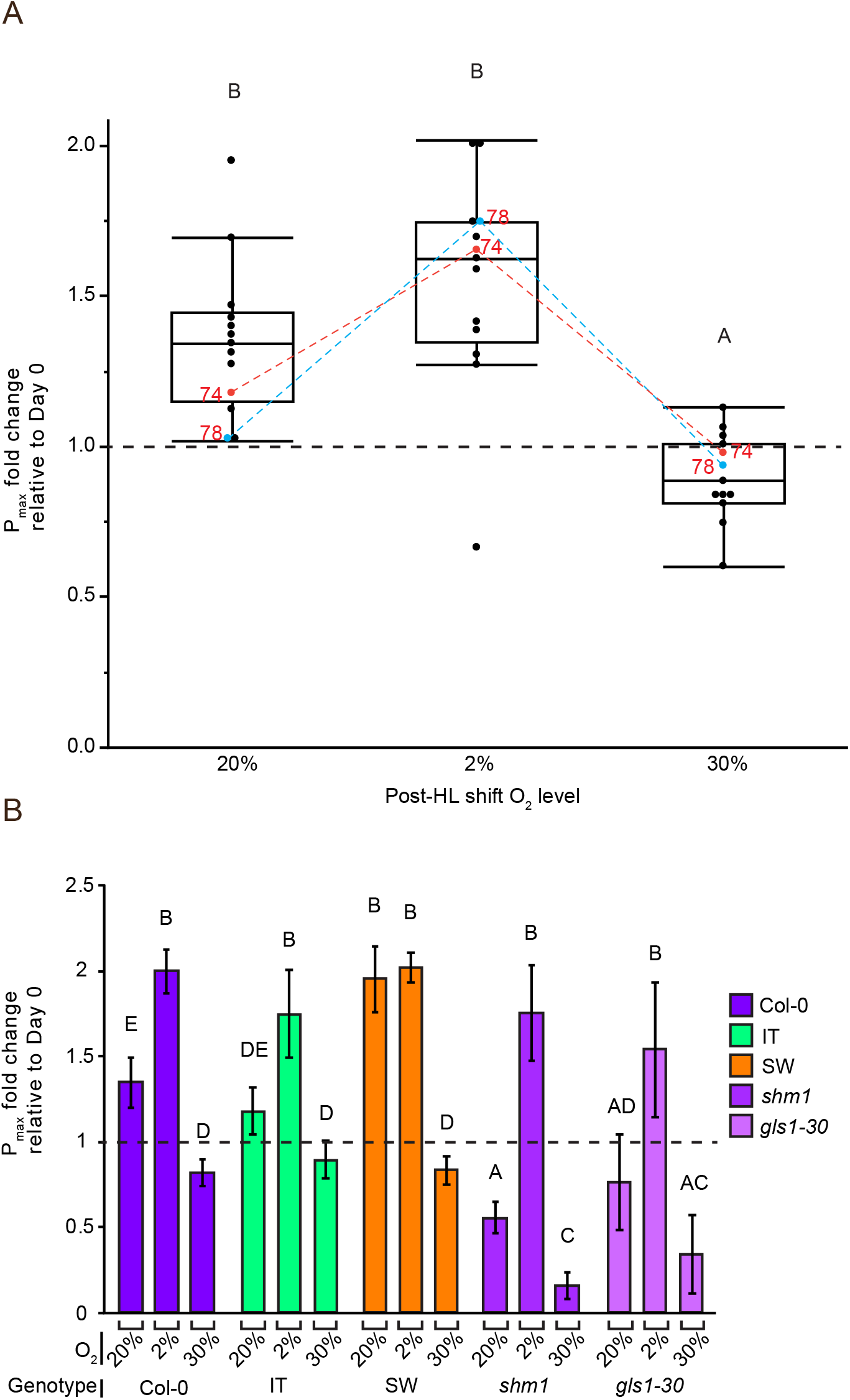
Low oxygen air enables photosynthetic capacity acclimation in HL in plants with weak responses in ambient oxygen conditions. (a) Photosynthetic capacity as measured by P_max_ per leaf area for Col-0, IT, SW, and a subset of ecotype selected from the larger accession panel(Cao et al., 2011) to capture the full phenotypic range of photosynthetic capacity acclimation responses were assayed at HL Day 4 (HL) in ambient O_2_ air, in low O_2_ air (400 ppm CO_2_, 2% O_2_, balance nitrogen), and in high O_2_ air (400 ppm CO_2_, 30% O_2_, balance nitrogen). Mean values for individual genotypes are shown as black circles for *n* = 3 biological replicates. (b) P_max_ at HL Day 4 and 7 in ambient O_2_ air, low O_2_ air, and high O_2_ air measured in Col-0 (purple), IT (green), SW (orange) ecotype, and two photorespiration mutants (serine-hydroxymethyltransferase 1 (*shm1*) and Fd-dependent glutamine-oxoglutarate glutamate aminotransferase 1-30 (*gls1-30*)), which were generated in the Col-0 background. Mean values ± standard errors (*n* = 3). Mean values that share the same letters are not statistically different, and those that do not share the same letters are statistically different based on one-way ANOVA and post hoc Tukey–Kramer HSD tests. Values for conditions where plants wilted and showed signs of chlorosis were not determined (ND).

In low O_2_ air (2% O_2_), the fold change in photosynthetic capacity at HL Day 4 was on average slightly greater than the fold change in ambient O_2_ (Fig. 8A). In contrast, high O_2_ air (30% O_2_) completely blocked the increase in photosynthetic capacity at HL Day 4. These results are inconsistent with the first model that high photorespiratory/N assimilation rates act as a positive signal necessary for the licensing of photosynthetic capacity acclimation in HL. Instead, this experiment strongly supports the second model in which low photorespiratory capacity represses photosynthetic capacity acclimation. Remarkably, non-acclimating and weakly acclimating accessions, such as Lago-1 (ecotype #79), Col-0, IT, Yeg-1 (ecotype #74), as well as the two photorespiratory mutants, were able to increase photosynthetic capacity specifically in low O_2_ air on HL Day 4 (Fig. 8A-B), a condition where photorespiration was repressed. Both Yeg-1 and Lago-1 are among the ecotypes with the lowest measured GS values on HL Day 1 (Fig. 7D), suggesting that, like the photorespiratory mutants, these ecotypes may have low photorespiratory capacity. Thus, insufficient photorespiratory capacity appears to have a repressive effect on the acclimation of photosynthetic capacity to HL, and certain weakly acclimating accessions are only able to increase photosynthetic capacity in HL under non-photorespiratory conditions.

Unlike in ambient O_2_ levels, photosynthetic capacity in HL did not continue to increase in low O_2_ at HL Day 7 (Fig. S11). In fact, measurements collected from Col-0, IT, and SW plants revealed that photosynthetic capacity was lower on HL Day 7 than on HL Day 4 in the low O_2_ grown plants (Fig. 8B, S11). This was an expected result because leaves developed in HL in low O_2_ conditions have low rates of N assimilation (Fig. 6) and may be unable long-term to increase investment in the synthesis of N rich photosynthetic machinery.

## Discussion

### Ecotypic variation in the acclimation of photosynthetic capacity to HL

The SW ecotype was shown to possess the traits of a strongly acclimating ecotype to HL relative to Col-0 and IT. The SW ecotype increased photosynthetic capacity (observable at HL Day 3) before Col-0 and IT and had the most oxidized Q_A_ redox state, as measured by q_L_, in the early phase of the HL shift (observable at HL Day 1) (Fig. 1, S2). Additionally, P_max_ per unit DW^−1^, a normalization that controls for differences in leaf thickness (Stewart et al., 2017; Athanasiou et al., 2010), attained its highest value in SW plants shifted into HL. This efficient response to HL fits with prior data suggesting that the SW ecotype (relative to the IT ecotype) is strongly HL adapted (Stewart et al., 2017).

While the timing of the induction of high photosynthetic capacity and the final P_max_ per unit DW differed between accessions, there were also shared patterns in the response of all three accessions to the shift in HL. The Q_A_ redox state began to recover (i.e., become more oxidized) on HL Day 1 in all three accessions (albeit most strongly in the SW ecotype), days prior to a measurable increase in photosynthetic capacity in any of the accessions (Fig. 1, S2). Further, all accessions eventually increased photosynthetic capacity on unit area basis in response to growth in HL, but for the Col-0 and IT plants this did not occur until HL Day 6 after the development of thicker HL acclimated leaves (as indicated by increased DW per unit area). The early increase in q_L_ before changes in photosynthetic capacity reflects both the faster kinetics of the acclimation of light-harvesting to HL growth (relative to the slower kinetics of the acclimation of photosynthetic capacity) and metabolic acclimation to increase flux through triose-phosphate utilization and N metabolism pathways, thus preventing feedback inhibition of PET (Demmig-Adams et al., 2014; Paul and Pellny, 2003; Pammenter et al., 1993). As discussed in greater detail below, the metabolic data collected on these plants supports that the early recovery of Q_A_ redox poise and photosynthetic capacity acclimation on HL Day 4 in the SW ecotype is tied to the ability of this ecotype to efficiently accelerate flux through triose-phosphate utilization and N metabolism pathways early in the HL shift relative to the IT and Col-0 plants.

A continuous distribution of photosynthetic capacity acclimation values (HL Day 4/LL) with fold changes of ∼0.95 to 2.2 was measured using the larger ecotype panel (Cao et al., 2011) (Fig. 2). While many accessions efficiently increased photosynthetic capacity by HL Day 4, the most strongly acclimating accessions did not geographically cluster. For instance, strongly acclimating ecotypes included the SW ecotype (latitude- 62°N), Dog-4 (Turkey, ecotype #5, latitude- 41.07°N), Star-8 (Germany, ecotype #25, latitude- 48.43°N), and Rovero-1 (Italy, ecotype #8, latitude- 46.25°N). This was not an unanticipated result. Ecophysiological studies of photosynthetic capacity acclimation to HL have suggested that adaptation to local growth environment is the major determinant of the strength of this response and in particular, plants that are dual-adapted to thrive in both shaded and full sunlight environments appear to have the largest dynamic range in photosynthetic capacity in response to light levels (Murchie and Horton, 1997; Björkman and Holmgren, 1963). Thus, we predict that the weakly acclimating accessions identified here represent shade-adapted *A. thaliana* ecotypes, whereas the strongly acclimating accessions are ecotypes able to thrive in full sunlight growth conditions and perhaps are also dual-adapted for growth in shade and full sun.

Parallels exist in the acclimation of photosynthetic capacity to cold and HL (Strand et al., 1999; Huner et al., 2012). There was a weak effect of latitude on photosynthetic capacity acclimation in this study (Table S1). The 12 northern latitude ecotypes (> 50°N) had a larger HL/LL fold change in photosynthetic capacity (1.75 for ETR_max_ and 1.57 for P_max_) relative to the 16 southern latitude ecotypes (< 40°N, 1.61 for ETR_max_ and 1.39). Exceptionally high photosynthetic capacity per unit area can be achieved by combining HL and cold growth conditions (Demmig-Adams et al., 2018; Cohu et al., 2013). It seems likely that latitudinal effects on photosynthetic capacity would be stronger using a combination of cold and HL during growth as opposed to treatment with only HL, the latter being the condition that most strongly favors the photoprotective contributions of photorespiration to HL acclimation (Savitch et al., 2000; Ma et al., 2014).

### Metabolic acclimation to HL growth conditions that increase photosynthetic capacity

Plants that increase photosynthetic capacity in HL growth will depend in turn on the capacity of metabolic sinks to utilize the products of enhanced PET rates. Here, we discuss differences in the strongly acclimating SW and weakly acclimating Col-0 in three metabolic pathways that support high PET rates by consuming products of PET (N assimilation, photorespiration, and triose-phosphate utilization) are explored in greater depth (Fig. 3-5). First, evidence for an increased rate of primary N assimilation in HL-shifted plants can be found in the increased abundance of Glu-derived AAs and the progressively increasing Gln/Glu ratio (Fig. 3). HL-shifted plants produce reducing equivalents in the light at an accelerated rate, and N assimilation acts as an important electron sink under these conditions (Foyer et al., 2003; Osmond et al., 1997). Total AA abundance increased over time in the HL-shifted plants, primarily driven by this change in the abundance of Gln and other Glu-derived AAs (Fig. 3). The abundance of Glu is tightly regulated in plants (Foyer et al., 2003; Huppe and Turpin, 1994; Florian et al., 2013), but Glu also increased on HL Day 6 and Day 10 in concert with the largest jumps in photosynthetic capacity measured on a leaf area basis.

Increased rates of N assimilation in HL-shifted plants depends on a constant supply of α-ketoglutarate carbon skeletons (Fig. S1). α-ketoglutarate levels, which constituted just 0.5% of the total OA pool, gradually declined in HL over the time course as photosynthetic capacity increased (Fig. 3). Early on in the HL shift, enhanced rates of *de novo* OA synthesis, particularly notable on HL Days 1 and 3, appear to have supported this α-ketoglutarate supply to N assimilation. PEP carboxylase activity, the key enzyme in *de novo* OA synthesis in plants, was increased on HL Day 1, and PEP abundance decreased in HL-shifted SW plants (Fig. 3-4, Fig. S6). The size of the total OA pool also increased slightly on HL Day 1 and 3, despite elevated AA synthesis rates (Fig. 3). In both Col-0 and SW, trehalose-6P was induced in HL-shifted plants. Trehalose-6P signals the repression of sucrose synthesis and enhances OA synthesis rates via activating PEP carboxylase activity (Figueroa et al., 2016). Consistent with this role for trehalose-6P, levels of sucrose biosynthesis intermediates (FBP, F6P, G6P, and Suc6P) dramatically increased in both accessions on HL Day 1 and cytosolic fructose bisphosphatase activity was reduced (Fig. 3, S6). Subsequently, FBP and Suc6P levels each declined on Day 3 and continued to decline for the rest of the HL growth time course (Fig. 3). The elevated FBP levels on Day 1 and Day 3 also may have allosterically activated pyruvate kinase, further contributing to the accelerated OA synthesis rates (Huppe and Turpin, 1994; Smith et al., 2000). While trehalose-6P acts to support accelerated OA synthesis rates in HL acclimating plants, weaker HL-acclimating genotypes often induce higher levels of trehalose-6P than stronger acclimating ones, as observed here by the stronger induction of trehalose-6P in Col-0 than in the SW ecotype (Fig. 3). Thus, a linear relationship between trehalose-6P levels and the induction of high photosynthetic capacity in HL remains unlikely (Paul, 2007; Dyson et al., 2015).

OA levels then dropped on HL Day 6 and then again on HL Day 10, driven predominantly by declining fumarate abundance and to a lesser extent isocitrate and α-ketoglutarate levels (Fig. 3,5). Thus, later in the HL shift, enhanced rates of *de novo* OA synthesis may have given way to the drawing down of the existing OA pool in the leaf as the primary mechanism to support enhanced α-ketoglutarate supply to N assimilation. The AA/OA ratio shifted from 0.88 in LL-grown plants to 3.33 on HL Day 10. In the extreme, the Gln/fumarate ratio was 0.28 in LL-grown plants (fumarate constituted 35% of the total OA pool in LL-grown plants) and increased to 13.14 by HL Day 10. High fumarate levels are necessary for the rapid assimilation of exogenous N in *A. thaliana* (Pracharoenwattana et al., 2010). Thus, we propose that a shift in the supply of α-ketoglutarate occurred in the SW ecotype across the HL time course. Prior to increase in photosynthetic capacity, accelerated flux through the anaplerotic reaction was primarily responsible for supplying carbon skeletons for N assimilation. However, once photosynthetic capacity began to increase, flux through the anaplerotic reactions appeared to decline as drawing down existing pools of OA, primarily fumarate, became the predominant source of α-ketoglutarate to maintain accelerated N assimilation rates.

High rates of photorespiration can also act to maintain Q_A_ redox poise in HL growth. The shift to HL in the Col-0, IT, and SW on HL Day 1 decreased *g_s_*, which in turn is likely to further accelerate Rubisco oxygenation rates (Fig. S7). The Gly/Ser ratio increased from 0.55 in LL-grown SW plants to 2.6 on HL Day 10, as a consequence of both progressively increasing Gly levels and declining Ser levels (Fig. 3). Photorespiratory and nitrate assimilation rates are linked (Bloom et al., 2002; Rachmilevitch et al., 2004; Busch et al., 2018), as exemplified in this work by the suppression of the HL-induced increase in Gly/Ser (a metabolic indicator of photorespiration) and Gln/Glu (a metabolic indicator of primary nitrogen assimilation) in low O_2_ (Fig. 6). One hypothesis for the mechanistic link between N assimilation and photorespiration rates is that excess sugar phosphates in HL are converted into the photorespiratory intermediates Gly and Ser, providing an additional N assimilation path under these conditions (Busch et al., 2018). However, under the conditions used in this work, which were selected to optimally induce increased photosynthetic capacity, the increase in Gln and Glu-derived AAs in HL-shifted plants far outpaced the increase in Gly levels (Fig. 3), indicating that high rates of N assimilation in HL-shifted was a direct consequence of primary nitrogen assimilation and not a secondary consequence of elevated photorespiratory-mediated AA synthesis.

The elevated synthesis of photosynthate in plants with increased photosynthetic capacity in HL will also depend on having sufficient triose-phosphate utilization capacity to keep pace with these enhanced synthesis rates. Sugar phosphates and sugars in Col-0 and SW plants, in particular on HL Day 1, both increased but to a lesser extent in the stronger acclimating SW ecotype (Fig. 3). Notably, the Fru + Glc level was induced 2.6-fold on HL Day 1 in SW plants and ∼1.5 fold on the subsequent sampling days, whereas Fru + Glc was 6.7-fold induced in Col-0 on HL Day 1. Sucrose, starch, CBB cycle, and glycolysis intermediates were all at higher levels on HL Day 1 in Col-0 relative to SW (Fig. 3-4). Furthermore, the increase in ATP/ADP ratio which occurs in the SW ecotype in HL was not observed in Col-0 (Fig. 3, 5). Each of these results points towards higher levels of sink limitation in the weakly acclimating Col-0 accession relative to the strongly acclimating SW ecotype following the HL shift (Demmig-Adams et al., 2017; Häusler et al., 2014; Goldschmidt and Huber, 1992; Paul and Pellny, 2003; von Schaewen et al., 1990). In summary, within the metabolomics data, it is possible to observe how the strongly acclimating SW ecotype efficiently accelerates rates of N assimilation, photorespiration, and sink utilization of photosynthate relative to Col-0, thereby, restoring redox poise to the PET in HL and allowing for the induction of increased in photosynthetic capacity in HL (Fig. 9).

**Figure 9.**
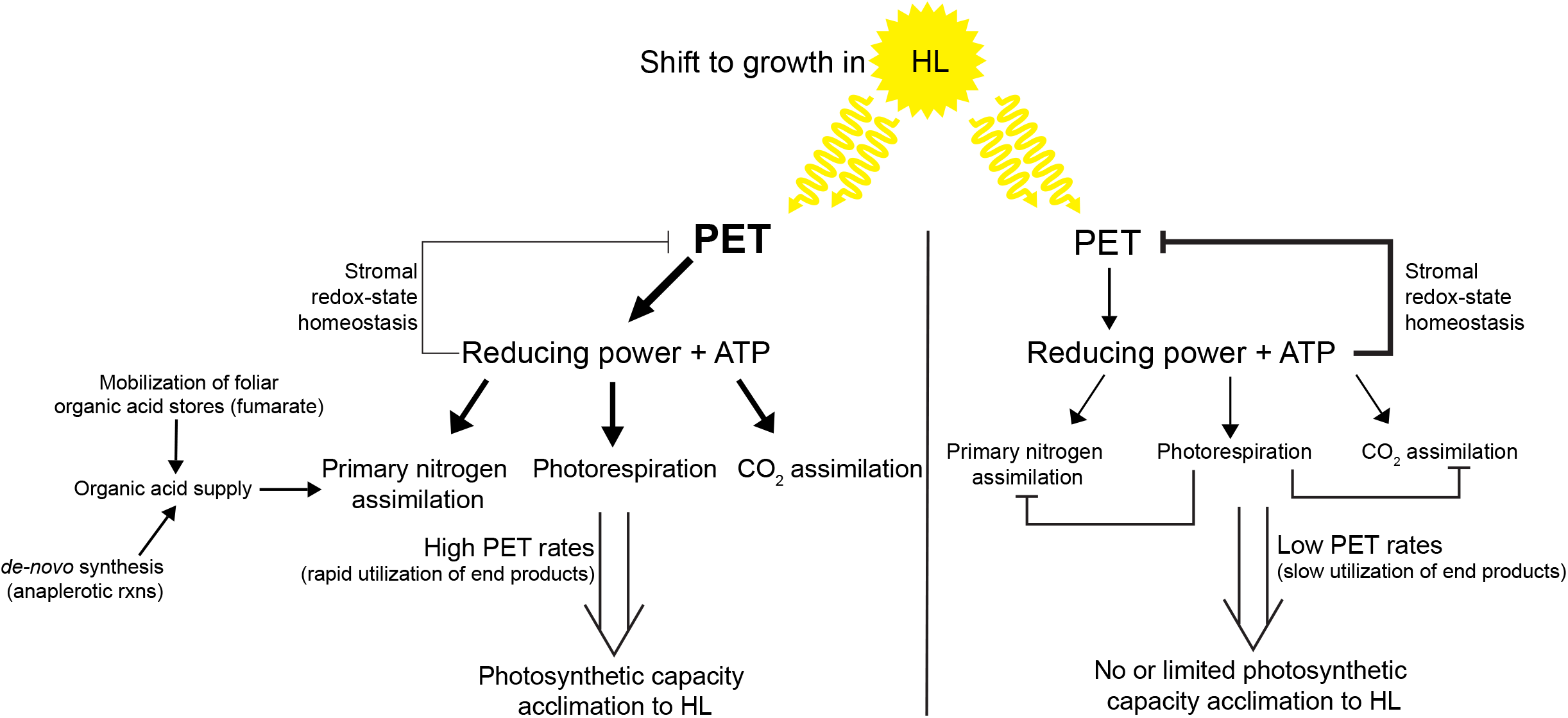
Model of the control of photosynthetic capacity acclimation by nitrogen assimilation and photorespiration pathways. The left side presents the situation for a genotype with strong photosynthetic capacity acclimation to HL. Rapid utilization of end-products of PET in the chloroplast via pathways including N assimilation and photorespiration maintains an oxidized Q_A_ state, limiting feedback inhibition and licensing photosynthetic capacity acclimation to HL. The right side presents the situation for a genotype with weak photosynthetic capacity acclimation to HL. Limited photorespiratory capacity represses the utilization of products of PET, in turn repressing PET rates.

### High N assimilation and photorespiration rates early in the HL shift correlate with the increase in photosynthetic capacity

The variation in photosynthetic capacity acclimation across the large *A. thaliana* accession panel provided an opportunity to test the links between metabolic differences and characteristics of photosynthetic capacity acclimation. Rates of N assimilation and photorespiration, as measured by Gln/Glu and Gly/Ser, respectively, on HL Day 1 were strongly correlated with the fold change in photosynthetic capacity on HL Day 4 measured as ETR_max_ (Fig. 7, Fig. S8-10). GS enzymatic activity also correlated, albeit not as strongly, with the fold change in photosynthetic capacity. Given the relationship between GS capacity to both photorespiration and N assimilation capacity (Kozaki and Takeba, 1996), this result suggests that N assimilation and photorespiration capacity were higher in plants that went on to increase photosynthetic capacity at HL Day 4. Further supporting that high photorespiratory capacity early in the HL shift is predictive of the subsequent induction of high photosynthetic capacity, the total Ser + Gly pool, largely driven by the increase in Gly, also correlated with the induction of photosynthetic capacity. As previously noted, N assimilation and photorespiration rates in HL are tightly linked (Bloom et al., 2002; Rachmilevitch et al., 2004; Busch et al., 2018), which again was reflected here in the shared behavior of the Gly/Ser and Gln/Glu levels across the *A. thaliana* accession population.

Low foliar C/N ratios have also been linked to higher photosynthetic capacity (Givnish, 1988; Evans and Poorter, 2001; Paul and Pellny, 2003). Here, the early induction of high N assimilation rates will allow plants that increase photosynthetic capacity at HL Day 4 to maintain a lower C/N ratio. This was also shown in the metabolomic data in which sugar and sugar-phosphate levels were higher and AAs lower in the weakly acclimating Col-0 relative to the strongly acclimating SW ecotype on HL Day 1, reflecting an imbalance in C/N in Col-0 at the beginning of the HL shift (Fig. 3).

N assimilation rates in HL are proportional to the rate of PET (Foyer et al., 2003). Given the strong correlation between Gln/Glu values and the magnitude of the induction of photosynthetic capacity (Fig. 7), it can be inferred that those accessions that most strongly induced photosynthetic capacity at HL Day 4 may have also been the accessions with the highest rates of PET under growth conditions on HL Day 1 (Fig. 9). This conclusion also fits with the observation that the SW ecotype had a more oxidized Q_A_ redox state early in the HL shift and had the highest Gln/Glu ratio relative to Col-0 and IT (Fig. 1, 6, S1). Thus, changes quite early in the HL shift in PET activity appear to set in motion a metabolic and signaling response that determines the magnitude of the induction of photosynthetic capacity that does not manifest until several days later in the HL time course.

### Repression of photosynthetic capacity acclimation in plants with low photorespiratory capacity

Two models can be envisioned for the signaling relationship between photorespiration and photosynthetic capacity acclimation to HL. The first is that high photorespiration rates act as a positive metabolic signal that directly initiates an increase in photosynthetic capacity. In this case, for instance, high N assimilation and photorespiration rates might initiate a signaling cascade that increases investment in photosynthetic machinery. The second, and not necessarily mutually exclusive, model is that low photorespiratory capacity generates a repressive signal that blocks photosynthetic capacity acclimation to HL. Consistent with this second model, the measurement of photosynthetic capacity acclimation to HL in low O_2_ air (a condition that represses the RuBP oxygenation in HL) was sufficient to convert weakly acclimating accessions into strongly acclimating ones (Fig. 8). Further, all strongly acclimating ecotypes tested were able to induce photosynthetic capacity to similar magnitudes in low O_2_ and ambient O_2_ on HL Day 4. High O_2_ air blocked the acclimation of photosynthetic capacity to HL in all plants tested, indicating that photorespiratory capacity can be overwhelmed in even the strongest acclimating ecotypes, thereby, blocking photosynthetic capacity acclimation.

Lago-1 (ecotype #79) was the most striking example of a non-acclimating ecotype under ambient air conditions that then strongly induced high photosynthetic capacity specifically in low O_2_ air (Fig. 8). Lago-1 behaved similarly to actual photorespiratory mutants (Somerville and Ogren, 1981, 1980), as characterized by non-acclimation of photosynthetic capacity in HL in ambient O_2_ conditions but strong induction in low O_2_ air (Fig. 8). The shared behavior of weakly or even non-acclimating ecotypes such as Lago-1 and photorespiratory mutants is compelling evidence that a characteristic of non-acclimating ecotypes may be low photorespiratory capacity. Additional support for this conclusion can be found here in the low GS enzyme activity levels, an indicator of both photorespiratory and N assimilatory capacity, low Gly/Ser, and low Gly + Ser levels on HL Day 1 in Lago-1 and other non-acclimating ecotypes (Figure 7). The weakly acclimating IT and Col-0 accessions behaved similarly to Lago-1 in low O_2_ air (Fig. 8), but in addition to the measurement of low GS enzyme activity in these plants, the PIB was also measured in these two genotypes (the PIB is proportional to the photorespiratory GDC activity). Again, pointing towards lower photorespiratory capacity in weakly acclimating accessions, the PIB was lower in Col-0 and IT before HL Day 4 relative to the strongly acclimating SW ecotype (Fig. S6).

The signal that blocks photosynthetic capacity acclimation in HL in response to insufficient photorespiratory capacity is not identified by this work but one possibility is the photorespiratory intermediate 2-phosphoglycolate (2-PG). 2-PG acts as a potent inhibitor of the RuBP-regeneration phase of the CBB cycle in photorespiration mutants, thereby simultaneously repressing both photorespiration and photosynthesis (Flügel et al., 2017; Li et al., 2019). 2-PG levels are homeostatically controlled as excess 2-PG shuts down its own synthesis, and thus, 2-PG abundance does not increase in HL (Li *et al*., 2019). One potential mechanistic explanation for the control of photosynthetic capacity acclimation in HL is that insufficient photorespiratory capacity in weakly acclimating ecotypes might cause the transient over-accumulation of 2-PG in HL, lowering photorespiration, photosynthesis, and PET rates (Fig. 9).

This relationship between photorespiration and photosynthetic capacity acclimation suggest that the absence of photosynthetic capacity acclimation response to HL in many shade-adapted ecotypes (Björkman and Holmgren, 1963; Murchie and Horton, 1997) may be a consequence of decreased N-partitioning to photorespiratory enzymes. Low initial photorespiratory capacity upon a shift to HL growth in shade-adapted ecotypes would prevent such ecotypes from efficiently increasing photorespiratory rates, thus leading to the transient accumulation of negative metabolic signals that repress PET in HL.

While photorespiration and enhanced N assimilation rates appear to be transiently dispensable for acclimation of photosynthetic capacity in HL (as measured on HL Day 4), leaves that developed in HL in low O_2_ (measured on HL Day 7) had lower photosynthetic capacity relative to leaves that developed in HL in ambient O_2_ (Fig. S11). This was an expected result, as it is difficult to envision how high photosynthetic capacity could be maintained long-term in HL with the low N assimilation rates in low O_2_ air since high photosynthetic capacity depends on increasing investment in N rich photosynthetic machinery (Evans and Poorter, 2001). Partial anoxia has also been shown to have long-term adverse effects on growth, including the repression of photosynthesis (Priestley *et al*., 1988; Kawasaki *et al*., 2015). That the long-term maintenance of high photosynthetic capacity in HL depends on enhanced rates of N assimilation in HL conforms to our expectations, however, it is more surprising that the licensing of photosynthetic capacity acclimation to HL appears to not depend on a positive metabolic signal linked to enhanced N assimilation rates under all conditions (Fig. 9).

In summary, multiple independent lines of evidence are presented in support of the conclusion that plants with low photorespiratory and N assimilation capacity on HL Day 1 experience a negative, photorespiration-dependent signal that interferes with the licensing of photosynthetic capacity acclimation to HL (Fig. 9). Markers of N assimilation and photorespiration rates were positively correlated with the magnitude of increased photosynthetic capacity on HL Day 4, reflecting the high rates of PET early in the HL shift in strongly acclimating ecotypes that are capable of efficiently accelerating flux through these pathways in HL (Fig. 7). High photorespiratory capacity prevents generation of the negative photorespiratory-dependent signal that interferes with photosynthetic capacity acclimation. Limited photorespiratory capacity that impairs a plant’s ability to acclimate to different growth light regimes may have detrimental effects on biomass productivity, which could be suppressed by the use of an engineered photorespiratory bypass (South et al., 2019). In their work, South et al. found that introduction of a photorespiratory bypass had positive effects on growth beyond modeled expectations (South et al., 2019), potentially reflecting the consequences of enhanced HL acclimation due to the absence of photorespiration-dependent feedback repression of PET in these engineered plants. Future work on photosynthetic capacity acclimation should continue to push towards a mechanistic understanding of how the regulatory links between the pathways of central metabolism control photosynthetic capacity in response to growth environment signals. These insights will inevitably open up new targets for the optimization of photosynthesis in crop species and provide a better predictive framework for how engineered pathways will integrate with central metabolism.

## Acknowledgments

We thank the BioAnalytical Facility at the University of North Texas for the support with mass spectrometry analysis, Olga Gaidarenko for the IR camera images and advice on IR-image analysis, Barbara Demmig-Adams and Jared J. Stewart for advice on the development of growth conditions for this work, and Setsuko Wakao and Masakazu Iwai for their comments on the manuscript. This work was supported by the Gordon and Betty Moore Foundation through Grant GBMF 2550.03 to the Life Sciences Research Foundation [to C.R.B]. K.K.N. is an investigator of the Howard Hughes Medical Institute.

## Supplemental Materials

### Supplemental Figure Legends

**Figure S1.**
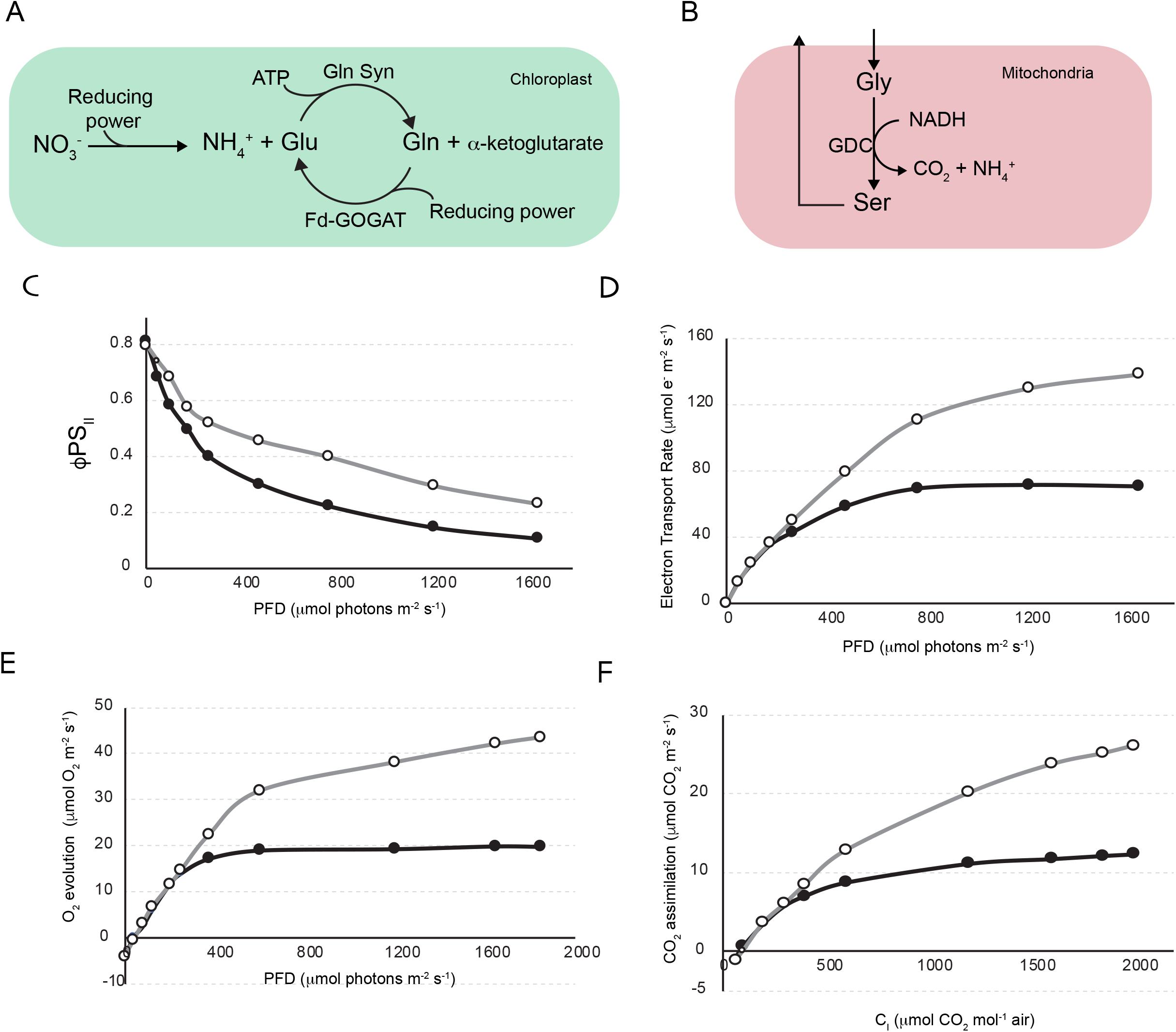
Determining conditions necessary for saturating light and CO_2_ to measure photosynthetic capacity and metabolic pathway depictions. (a) Simple cartoon of nitrogen assimilation in the chloroplast in plants. (b) Simple cartoon of glycine decarboxylase (GDC) activity in the mitochondria within the photorespiration pathway. (c-f) SW ecotype plants grown in LL (grey) or Day 10 after the shift to HL (black) assayed for (c) ΦPS_II_, (d) electron transport rate (μmol e^−^ m^−2^ s^−1^), and (e) O_2_ evolution rate (μmol e^−^ m^−2^ s^−1^) over a range of photon flux densities (PFD, μmol quanta m^−2^ s^−1^), and (f) CO_2_ net assimilation rate at saturating light over a range of intracellular CO_2_ concentrations (C_i_, μmol CO_2_ mol^−1^ air).

**Figure S2.**
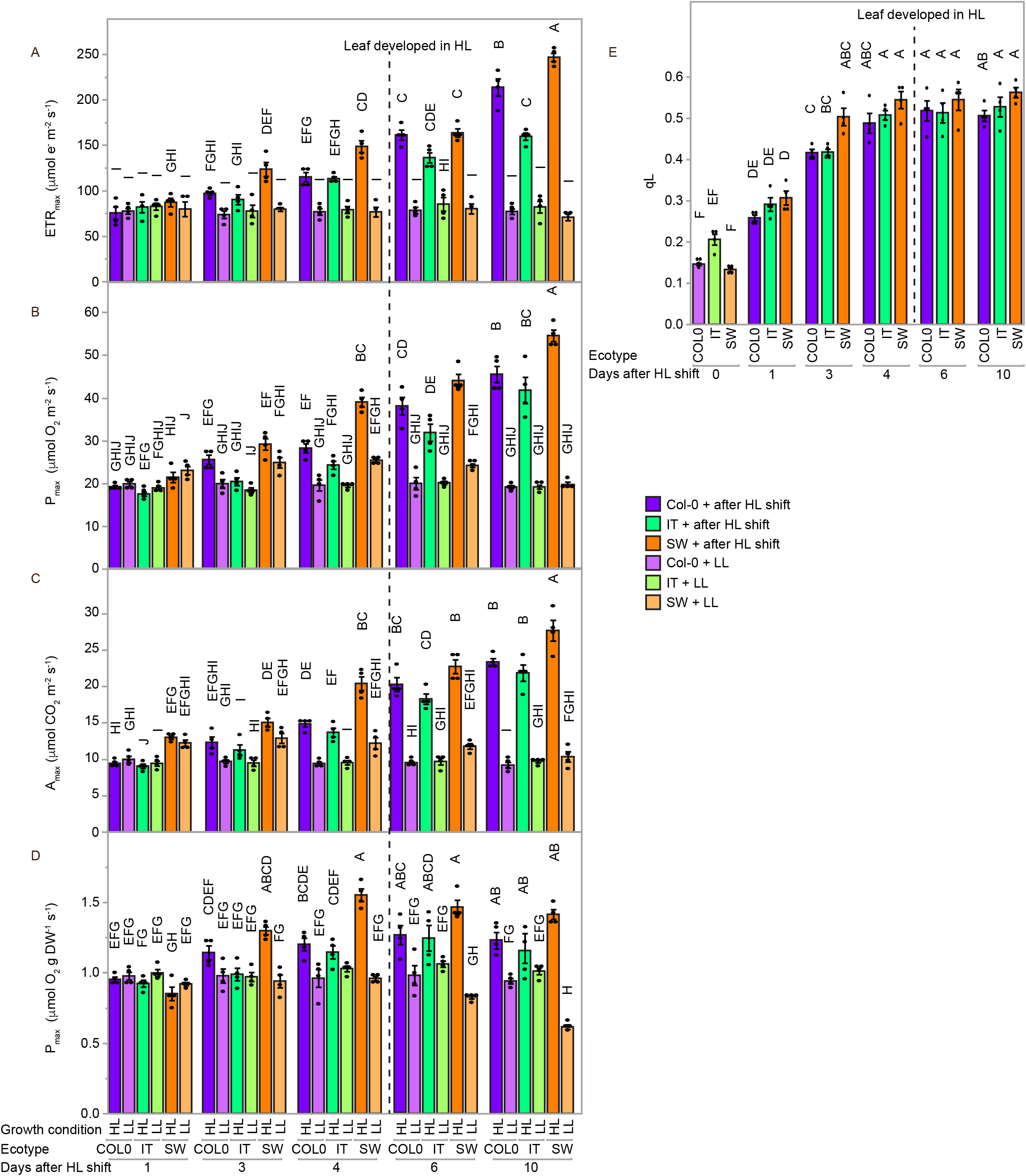
The dynamics of photosynthetic capacity acclimation to HL in the Col-0, IT, and SW plants. (a-d) Photosynthetic capacity as measured by (a) maximum electron transport rate, ETR_max_ (i.e., maximal light-saturated photosynthetic electron transport rate) per leaf area, (b) maximum oxygen evolution rate, P_max_ (i.e., maximal CO_2_- and light-saturated oxygen evolution rate) rate per area, (c) maximum carbon assimilation rate, A_max_ (i.e., maximal CO_2_- and light-saturated CO_2_ assimilation rate) per leaf area, and (d) P_max_ per unit dry leaf mass. (e) Q_A_ redox state as measured by q_L_ at growth light intensity. Col-0 (purple), IT (green), and SW (orange) plants that were grown in either low light (LL) or shifted to high-light (HL) at measurement Day 0. Mean values ± standard errors (*n* = 4) with individual data points displayed as dots (black). Mean values that share the same letters are not statistically different, and those that do not share the same letters are statistically different based on one-way ANOVA and post hoc Tukey–Kramer HSD tests.

**Figure S3.**
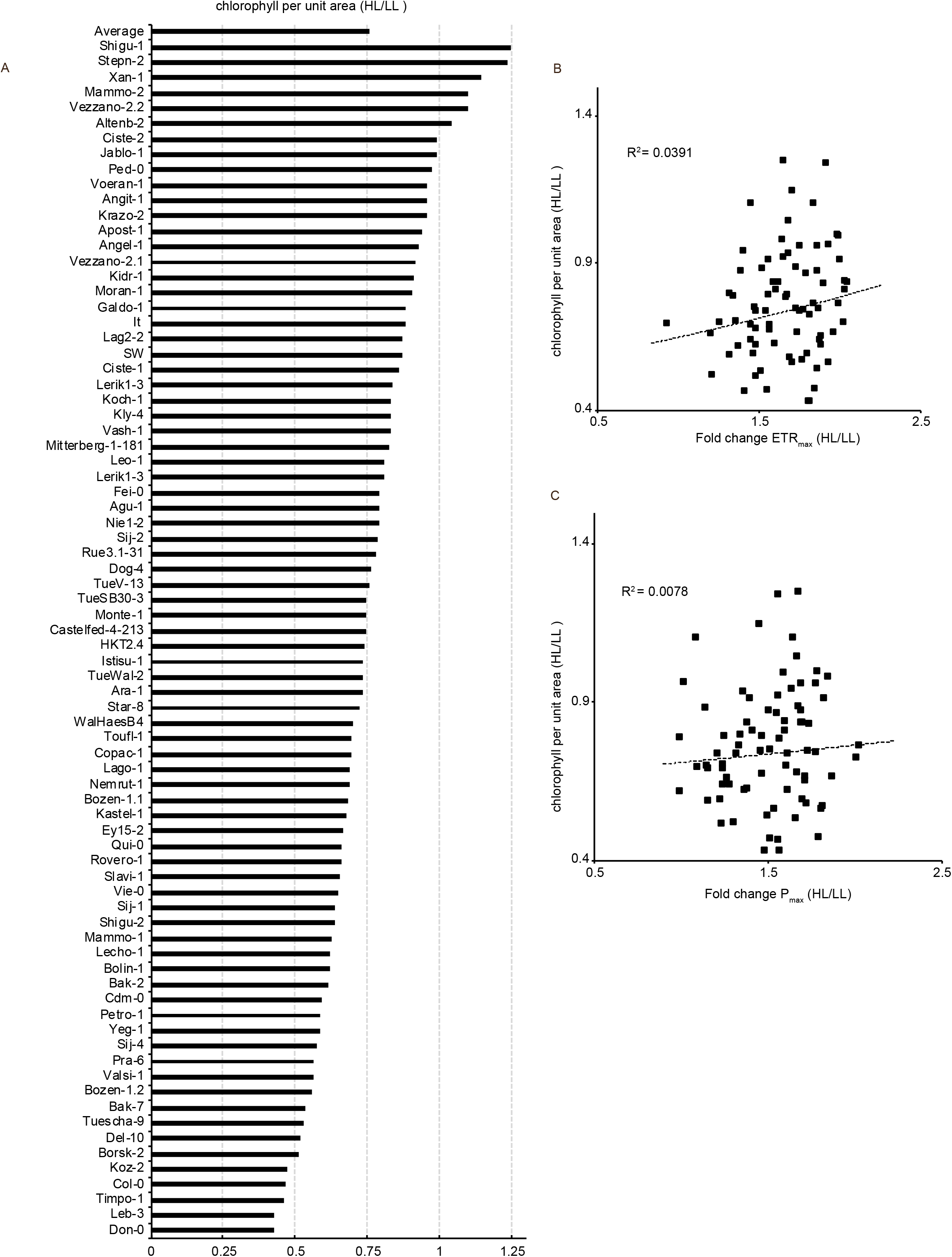
The response of chlorophyll per unit area to HL shift. (a) The ratio of chlorophyll per unit area in Day 4 HL-shifted (HL) over LL-grown (LL) plants for Col-0, IT, SW, and the accession panel (Cao et al., 2011). Mean values are calculated using *n* = 2 to 4. Scatter plots of fold change in chlorophyll per unit area (HL/LL) plotted against the fold change in photosynthetic capacity (HL/LL) as measured by (b) ETR_max_ and (c) P_max_ for the same plants as those used to quantify chlorophyll per unit area.

**Figure S4.**
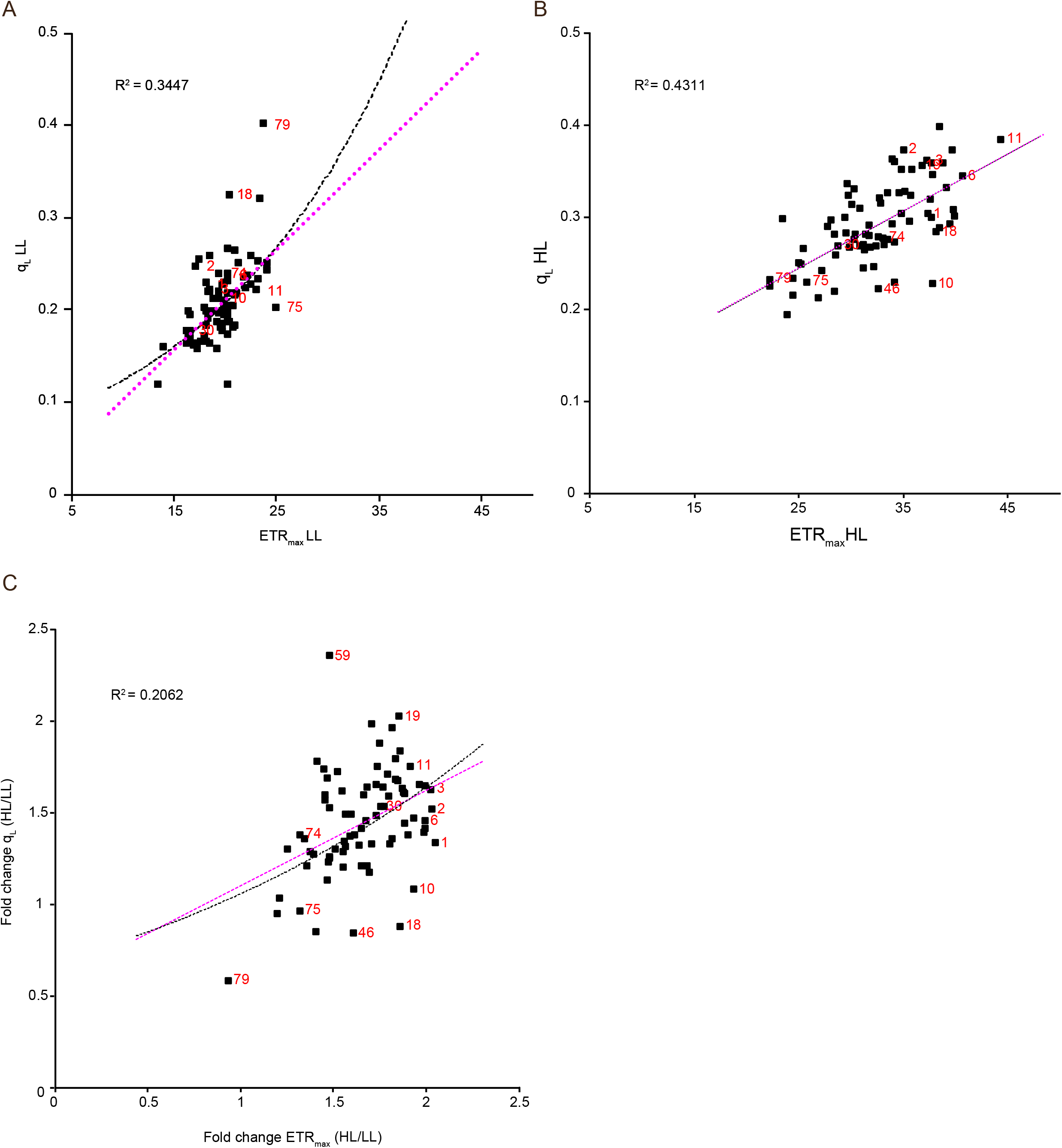
Natural diversity in Q_A_ redox state in response to an HL shift measured under saturating light. Correlation of Q_A_ redox state as measured by q_L_ with photosynthetic capacity as measured by ETR_max_ for Col-0, IT, SW, and the accession panel (Cao et al., 2011). (a) q_L_ and ETR_max_ assayed in LL-grown plants, (b) q_L_ and ETR_max_ assayed in Day 4 HL-shifted plants (HL), and (c) fold change (HL/LL) for q_L_ and ETR_max_. Best-fit regression and associated R^2^-values (black) and theoretical 1:1 regression (magenta) are displayed. Ecotype numbers (red) corresponding to the unique number for each ecotype Fig. 2A. ETR_max_ and q_L_ mean values were calculated using *n* = 2 to 4.

**Figure S5.**
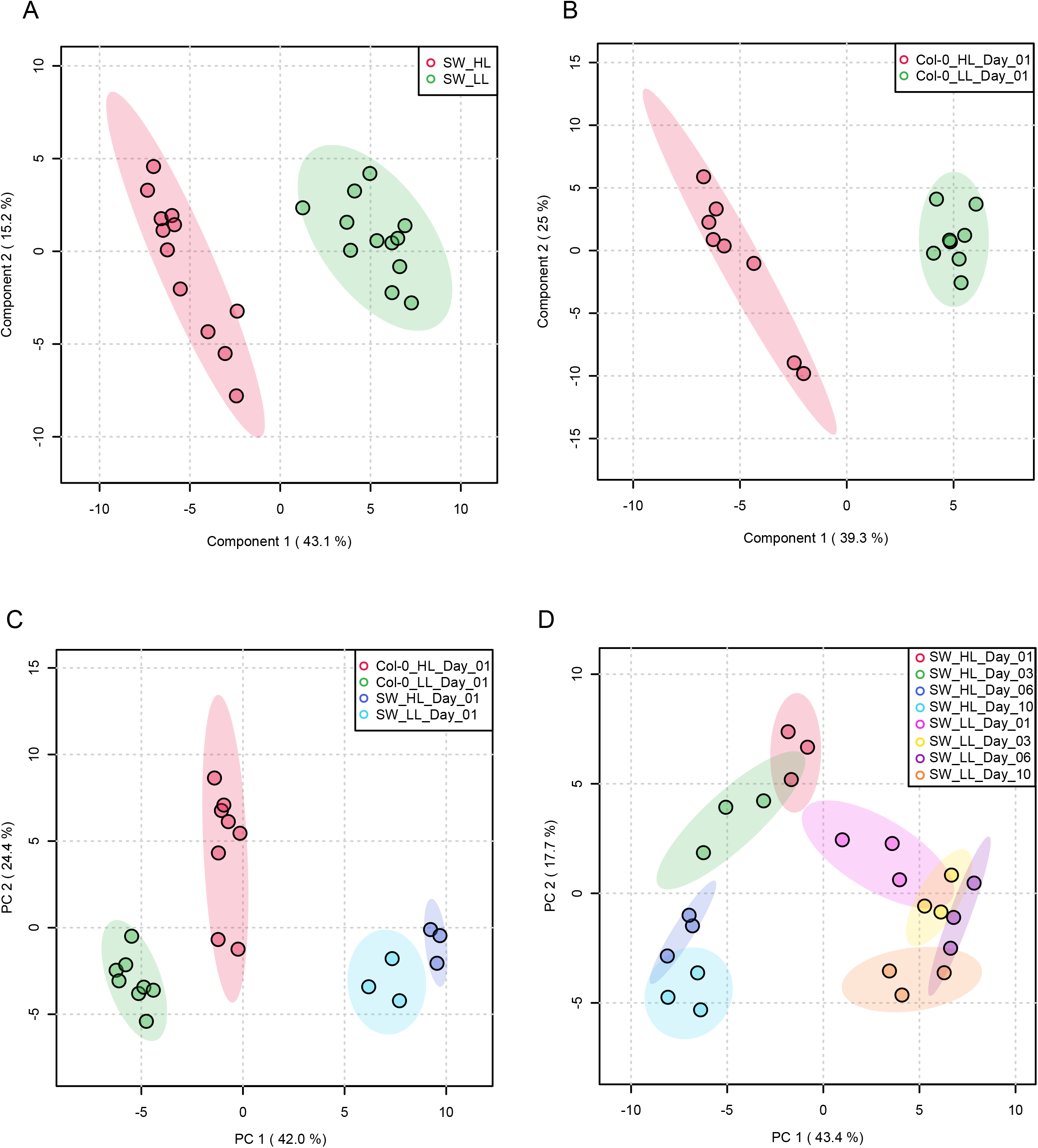
Differentiation of metabolomes by genotype, growth condition, and time component. Unsupervised principal component analysis (PCA) of metabolomic data from LL plants (green) and HL-shifted plants (pink) for the (a) SW and (b) Col-0 plants. The shaded regions in the PCA plots represent 95% confidence intervals. Unsupervised PCAs of metabolomic data for (c) SW LL (cyan), Col-0 LL (green), SW post-HL shift Day 1 (blue), and Col-0 post-HL Day 1 (red) plants, and (c) SW plants post-HL shift Day 10 (cyan), HL Day 6 (blue), HL Day 3 (green), HL Day 1 (red), LL grown plant Day 10 (orange), LL Day 6 (yellow), LL grown Day 3 (dark purple), and LL grown Day 1 (fuchsia).

**Figure S6.**
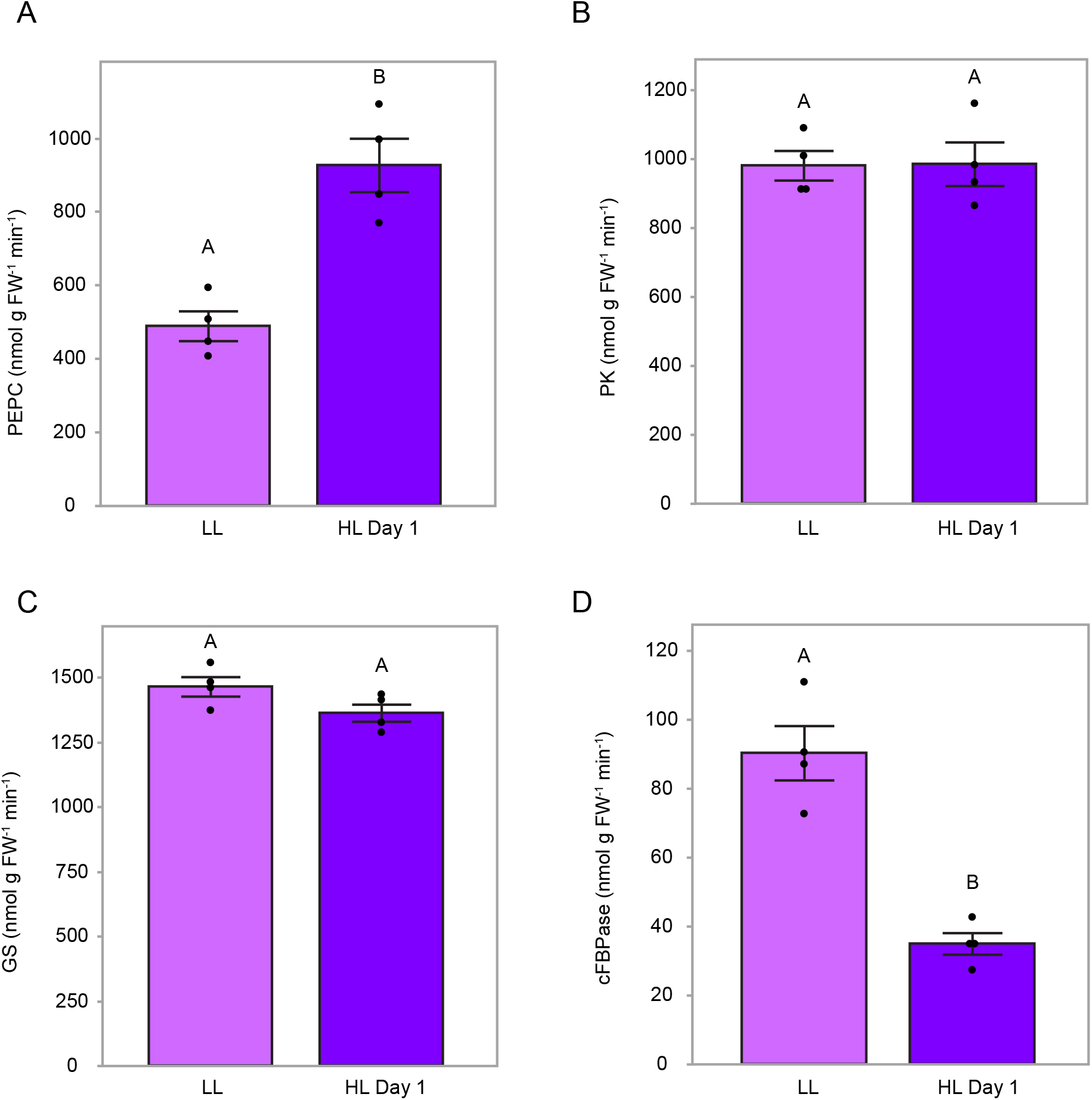
Enzyme activities in LL and HL-shifted plants. (a) PEP carboxylase (PEPC), (b) pyruvate kinase (PK), (c) glutamine synthetase (GS), and (d) cytosolic fructose bisphosphatase (cFBPase) enzymatic activities from leaf extracts for LL-grown (light purple) and HL Day 1 (purple) Col-0 plants. Mean values ± standard errors (*n* = 4). Mean values that share the same letters are not statistically different, and those that do not share the same letters are statistically different based on one-way ANOVA and post hoc Tukey–Kramer HSD tests.

**Figure S7.**
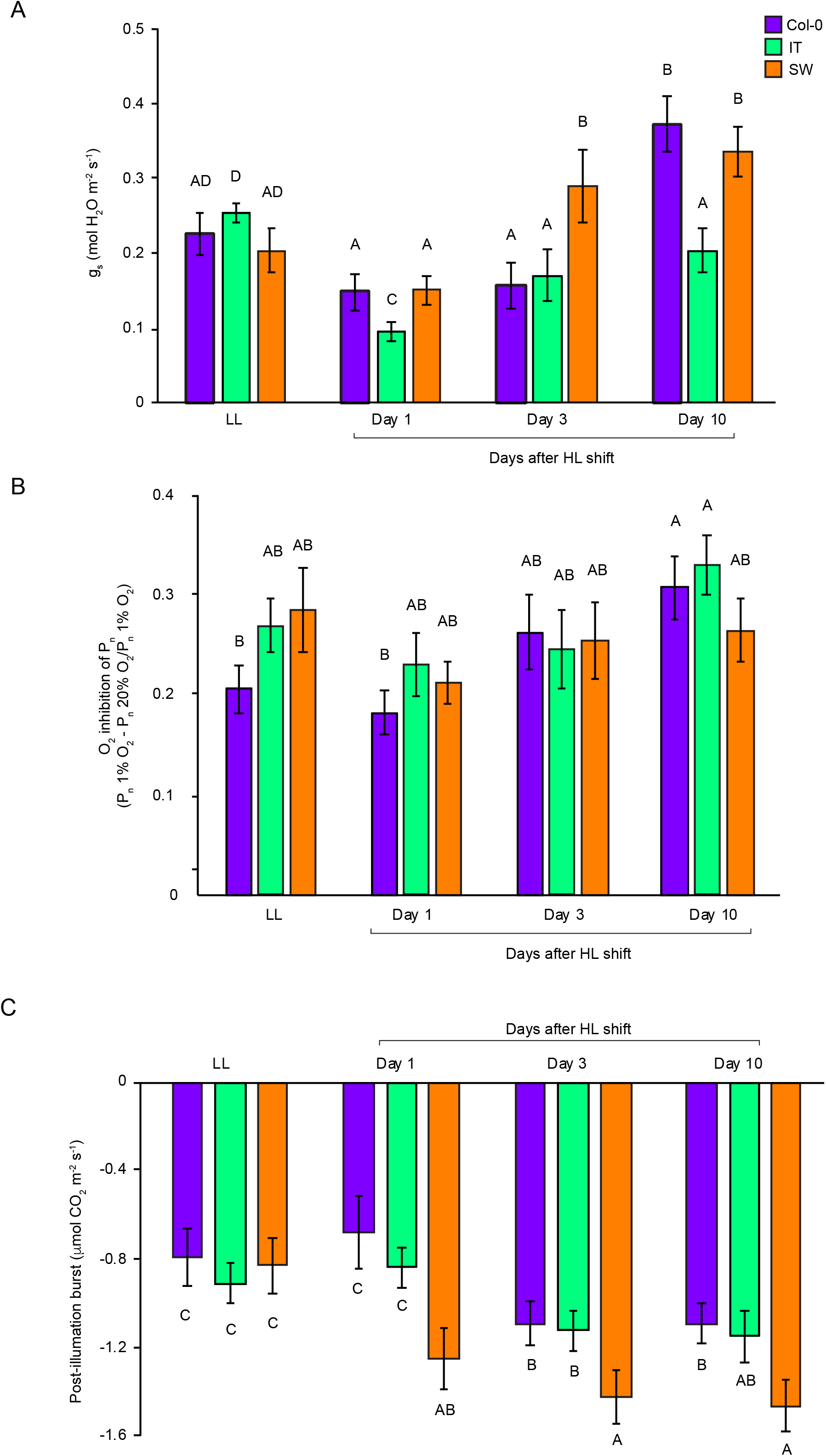
Response of gas exchange parameters related to photorespiration after the HL shift in Col-0, IT, and SW plants. (a) Stomatal conductance (mol H_2_0 m^−2^ s^−1^) measured under growth conditions, (b) O_2_ inhibition of net photosynthetic rate (P_n_) measured at saturating light, and (c) post-illumination CO_2_ burst (PIB) quantified as maximum respiration rate (μmol CO_2_ m^−2^ s^−1^) in the dark averaged over 10 seconds after 20 min of saturating light. Col-0 (purple), IT (green), and SW (orange) plants were grown in either low light (LL) or shifted to high-light (HL) at measurement Day 0. Mean values ± standard errors (*n* = 4). Mean values that share the same letters are not statistically different, and those that do not share the same letters are statistically different based on one-way ANOVA and post hoc Tukey–Kramer HSD tests.

**Figure S8.**
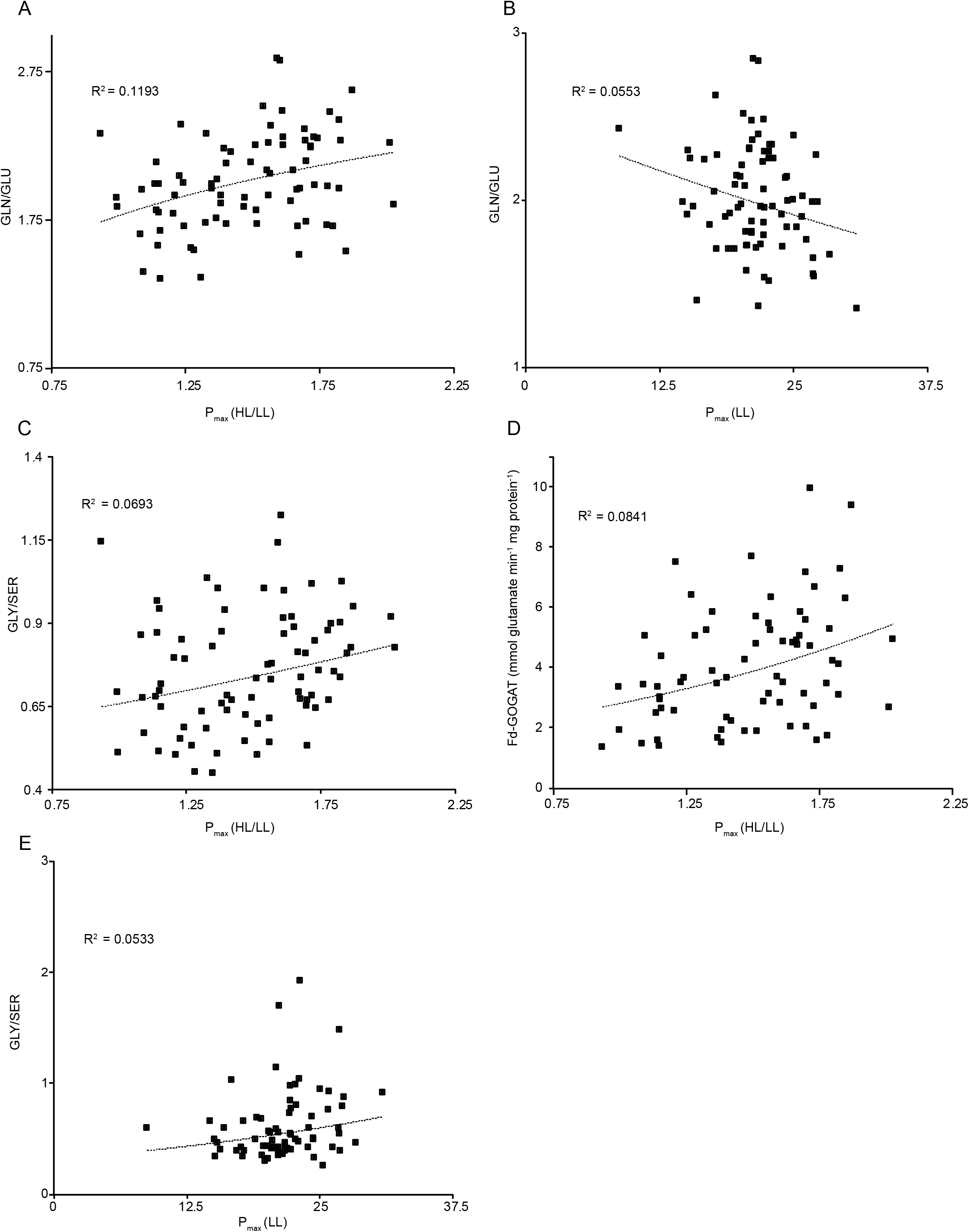
Correlation of metabolic indicators of photorespiration and nitrogen assimilation rates on HL Day 1 with increased photosynthetic capacity on HL Day 4 as measured P_max_. Correlation of Col-0, IT, SW, and the accession panel (Cao et al., 2011) for (a-c) fold change (HL/LL) in P_max_ in Day 4 post-HL-shift (HL) over LL-grown (LL) plants with (a) Gln/Glu ratio, (b) Gly/Ser ratio and (c) Fd-GOGAT (mmol Glu min^−1^ mg protein^−1^) on Day 1 of the post-HL shift. (d) Anticorrelation of P_max_ in LL grown plants with Gln/Glu on Day 1 after HL shift. (e) Correlation of P_max_ in LL grown plants with Gly/Ser ratio on Day 1 after HL shift. P_max_ mean values were calculated using *n* = 2 to 4. Enzymatic rate quantifications were calculated using *n* = 2 to 4 biological replicates and *n* = 2 technical replicates of each biological replicate. Ecotype numbers (red) that correspond to unique numbers in Fig. 2A are included.

**Figure S9.**
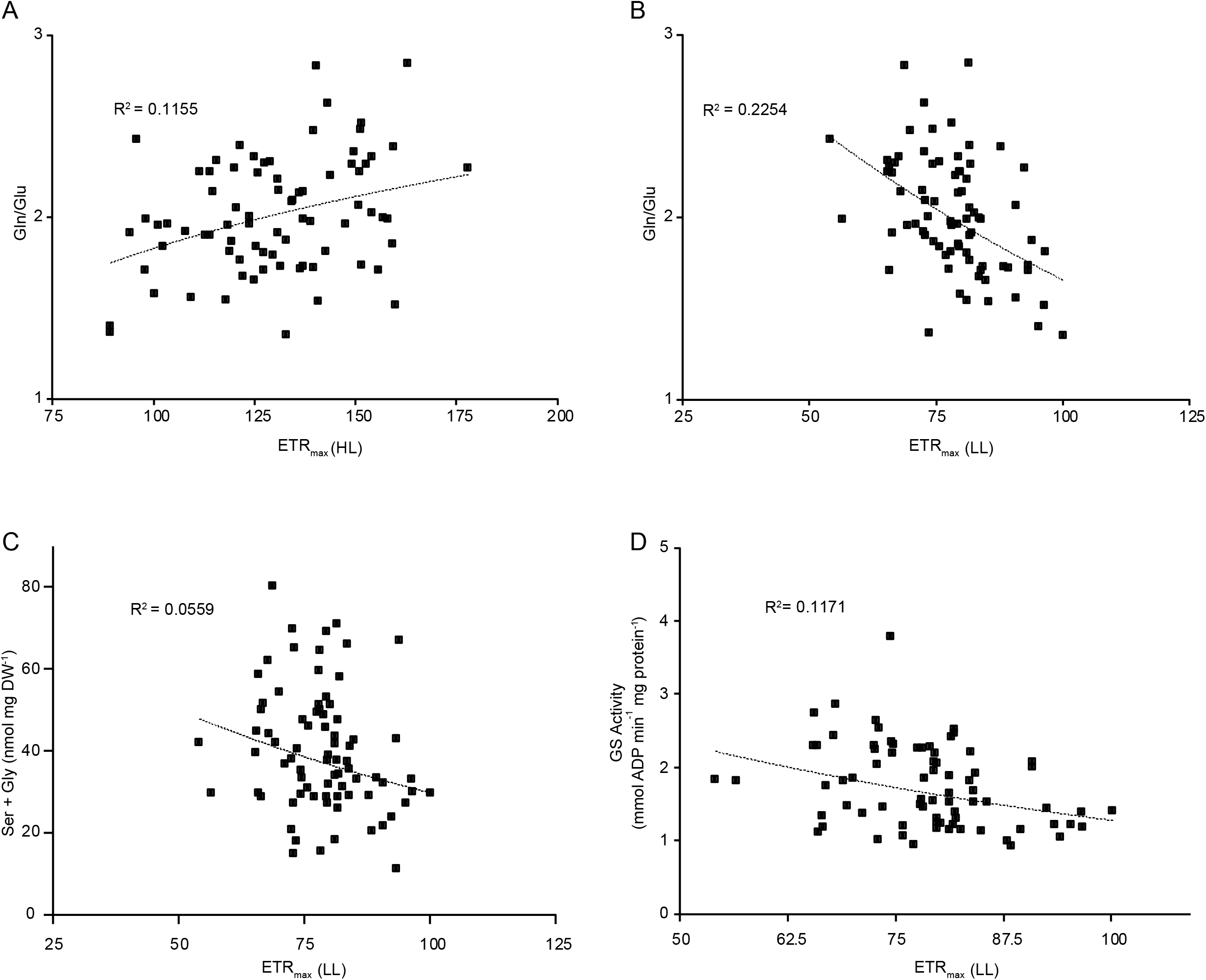
Correlation of metabolic indicators of photorespiration and nitrogen assimilation rates on HL Day 1 with increased photosynthetic capacity on HL Day 4 as measured by ETR_max_. Photosynthetic capacity as measured by ETR_max_ measured in (a) Day 4 after HL shift and (b-d) LL-grown plants for Col-0, IT, SW, and the accession panel (Cao et al., 2011) correlated with quantification of (a) Gln/Glu ratio and anticorrelated with (b) Gln/Glu ratio, (c) Gly + Ser (nmol mg DW^−1^), and (d) GS activity (mmol ADP min^−1^ mg protein^−1^) on Day 1 after HL shift. ETR_max_ mean values were calculated using *n* = 2 to 4. Enzymatic rates were calculated using *n* = 2 to 4 biological replicates and *n* = 2 technical replicates of each biological replicate. Ecotype numbers (red) that correspond to unique numbers in Fig. 2A are included.

**Figure S10.**
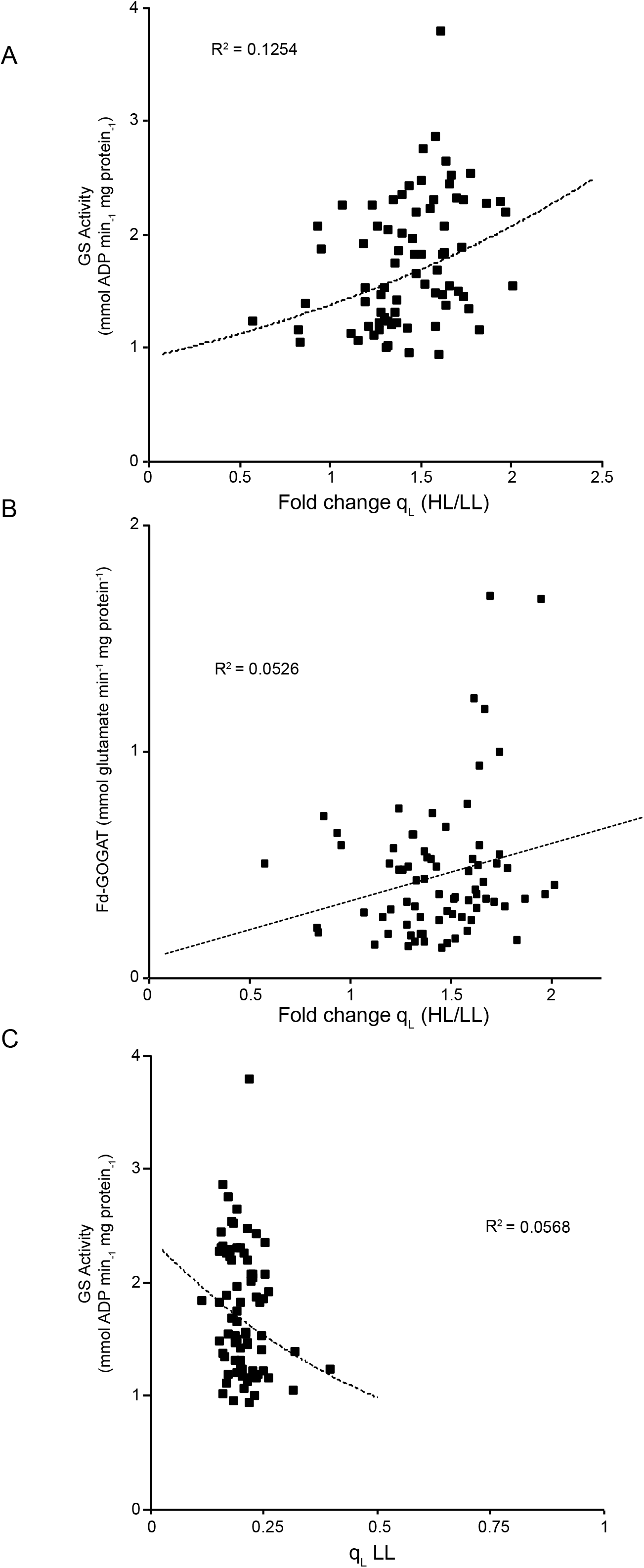
Correlation of metabolic indicators of photorespiration and nitrogen assimilation rates on HL Day 1 Q_A_ redox on HL Day 4 as measured in saturating light. Fold change (HL/LL) of the Q_A_ redox state as measured by q_L_ for Col-0, IT, SW, and the accession panel (Cao et al., 2011) correlated with quantification of (a) GS activity (mmol ADP min^−1^ mg protein^−1^), (b) Fd-GOGAT activity (mmol Glu min^−1^ mg protein^−1^) on Day 1 of the post-HL shift. (c) q_L_ from LL-grown plants anticorrelated with GS activity (mmol ADP min^−1^ mg protein^−1^) on Day 1 after HL shift. q_L_ mean values were calculated using *n* = 2 to 4. Enzymatic rate quantifications were calculated using *n* = 2 to 4 biological replicates and *n* = 2 technical replicates of each biological replicate.

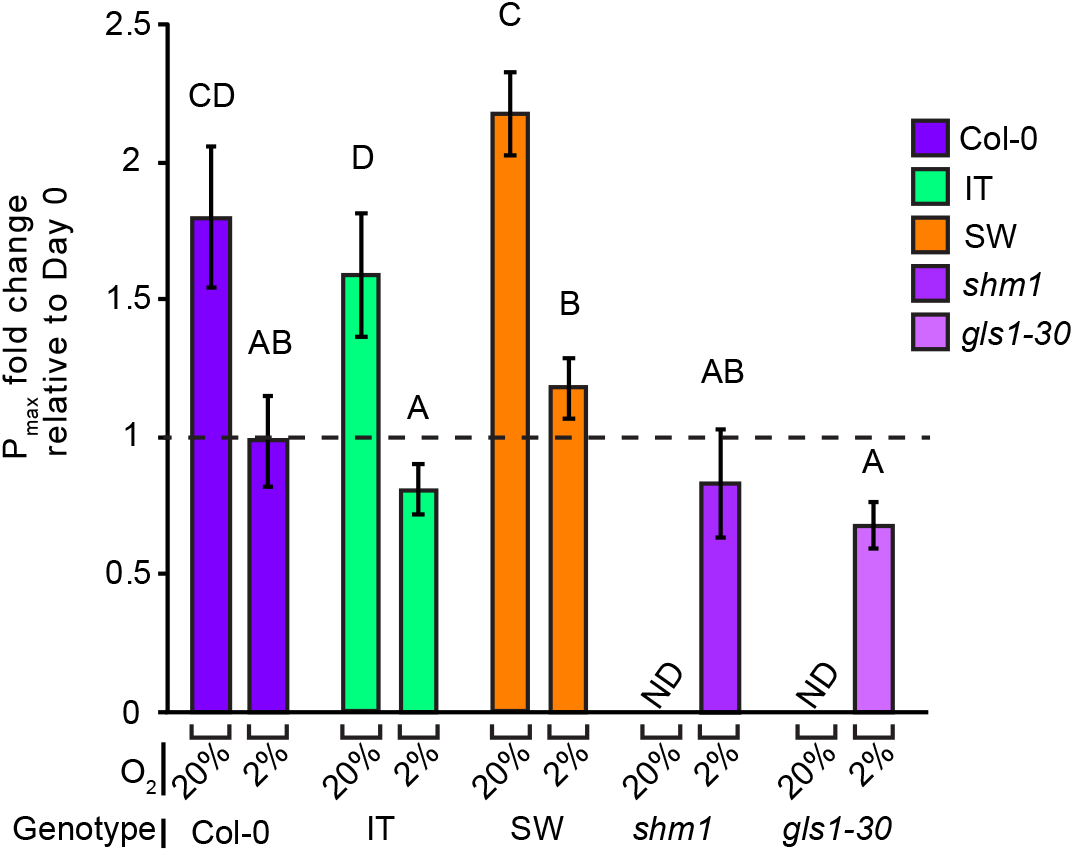

### Supplemental Tables

**Table S1.** Photosynthetic capacity values as measured in P_max_ and ETR_max_ and Q_A_ redox state as quantified by q_L_.

**Table S2.** LC-MS/MS metabolomics data for all conditions.

**Table S3.** Gln, Glu, Ser, Gly, GS activity, and Fd-GOGAT activity levels in Col-0, IT, SW, and the accession panel from Day 1 after HL shift.

